# Decoding brain anatomy from neuronal neighborhoods with MYCEL

**DOI:** 10.64898/2026.09.26.754562

**Authors:** Jesus Gonzalez-Ferrer, Avelina Moreno-Ochando, John R. Minnick, Aidan Schneider, Mircea Teodorescu, David Haussler, Mohammed A. Mostajo-Radji

**Affiliations:** Genomics Institute, University of California Santa Cruz, Santa Cruz, CA, 95064, United States; Live Cell Biotechnology Discovery Lab, University of California Santa Cruz, Santa Cruz, CA, 95064, United States; Department of Biomolecular Engineering, University of California Santa Cruz, Santa Cruz, CA, 95064, United States; Department of Statistics & Data Science, Yale University, New Haven, CT, 06511, United States; Department of Electrical and Computer Engineering, University of California Santa Cruz, Santa Cruz, CA, 95064, United States

## Abstract

Extracellular recordings carry information about neuronal location, and pooling per-unit predictions across neighboring electrodes improves its readout, but whether this local pooling should be learned is untested. We present MYCEL (Message-passing Yields Cellular Embeddings of Location), a graph neural network whose nodes are spike-sorted units carrying waveform and spike-timing features selected on cell-type ground truth, and whose edges encode recording geometry. On the Allen Brain Observatory Visual Coding Neuropixels dataset, learned message passing outperforms fixed aggregation, with the largest gains where single units are least informative. On the International Brain Laboratory Brain Wide Map, MYCEL matches the strongest population model at one-tenth the pairwise interactions. On held-out animals, the embeddings recover laminar depth within visual cortex. Retrained with only the edge geometry changed, the same framework decodes anatomy in macaque visual cortex and human medial temporal lobe. Thus, how neighboring neurons are combined matters as much as the features that describe them.

## Introduction

A central goal of systems neuroscience is to understand how the properties of individual neurons relate to the organization of the circuits and anatomical structures in which they reside^1^. Extracellular electrophysiology provides direct access to neuronal activity, and modern high-density probes can record hundreds to thousands of units simultaneously across multiple brain regions^2^. Yet the anatomical identity of those units is usually established independently of their activity, for example by reconstructing a probe trajectory and registering it to a reference atlas after the experiment^3,4^. As recordings become larger and extend across species and preparations, this separation raises the question of how much information about a neuron’s location can be recovered from its electrophysiology alone.

Individual neurons carry some of that information. Waveform shape, firing rate, burst structure and spike timing vary across neuronal classes and anatomical structures^5,6^, and methods for extracellular cell typing increasingly use richer combinations of waveform and spike-train structure^7–11^. A single unit’s electrophysiology therefore provides partial evidence about its location, and the units recorded alongside it appear to carry additional evidence^6^. One way to use the surrounding population is to learn neuronal embeddings from the activity of many simultaneously recorded units. Published models learn from population dynamics^12,13^, from contrastively trained spatiotemporal transformers over population spike trains^14^, or from spike-level tokenization across multisession populations ^15^. These models show that population activity carries substantial information about neuronal and anatomical identity. Local spatial context has also been used by pooling per-unit predictions across neighboring electrodes with a fixed rule^6,9^. Whether pooling over such a local, geometry-defined neighborhood should instead be learned has not been tested.

Here we introduce MYCEL (Message-passing Yields Cellular Embeddings of Location), which represents simultaneously recorded neurons as a spatial graph. Each node begins with intrinsic extracellular features describing waveform shape and spike timing, while edges connect nearby units according to the geometry of the recording. Message passing keeps a representation for every neuron, so the same graph can be read out for individual units, groups of neighboring units or the whole recording. To keep the node features interpretable and independent of the anatomical task, we search combinations of spike-derived features on ground-truth cell-type datasets, use the result to choose one fixed node representation, and do not tune it for brain region.

We evaluate MYCEL on mouse recordings from the Allen Brain Observatory Visual Coding Neuropixels dataset^4,16^ and the International Brain Laboratory (IBL) Brain Wide Map^17^. Individual-unit features carry above-chance anatomical information, and composing them across local populations substantially improves region decoding. Learned pooling improves further on fixed pooling over the same neighborhood, most on the finer Allen partitions. On the Brain Wide Map, MYCEL matches the strongest population model at a fraction of the training cost. The embeddings also retain anatomical information below the level of discrete region labels, recovering continuous position along cortical depth and, less certainly, along the hippocampal long axis. Finally, the same framework extends from mouse Neuropixels to macaque cortex, planar neural-organoid arrays and human medial temporal lobe recordings. Together, these results show that extracellular electrophysiology carries anatomical information across multiple spatial scales, and that reading each neuron together with its neighbors is an effective way to recover it.

## Results

### MYCEL composes cell-type features over a local graph

Methods for decoding cell type or brain region from extracellular spikes typically use hand-designed input representations, confounding the effects of the representation with those of the decoding model. To separate the two, we selected the per-unit features with a data-driven search. A non-dominated sorting genetic algorithm searched a library of spike-derived primitives, scoring each candidate with a conditional variational autoencoder (cVAE) evaluator held fixed across candidates, on three opto-tagged and expertly annotated cell-type datasets spanning two species and different recording technologies (Methods; Supplementary Figs. 1 and 2). The discovered representations statistically match or exceed the published fixed-encoding baselines on all three datasets. On Lisberger the best candidate reaches 0.78 against 0.70 for the strongest baseline (Welch’s *t* = 3.20, *P* = 0.014, Cohen’s *d* = 2.02, *n* = 5 folds each), and on Hull (0.93) and Buzsáki (0.63) it matches the published baselines^7,9–11^. The best representation differed between datasets, so no single hand-picked encoding is best everywhere (Methods). Autocorrelogram-derived scalars are the only primitive family selected on all three, though a different single primitive ranks highest on each (Supplementary Figs. 3 and 4). Because no single top candidate suits all three, we assembled one per-unit representation around the autocorrelogram scalars and carry it unchanged into every result below. We call these the *shape-and-timing features*. They consist of 57 scalars summarizing waveform shape, interspike intervals (ISIs), autocorrelogram structure and spectral power, concatenated with the 100-bin autocorrelogram histogram itself (157 dimensions per unit; Supplementary Fig. 5).

MYCEL combines the shape-and-timing features across a local population. Each sorted unit is a node carrying its shape-and-timing features, edges join units within a 200 *µ*m radius on the same probe, and a two-layer graph attention network (GATv2) ^18^ message-passes the features into a region call for every unit (Fig. 1a). Only the edge relation changes between preparations, so the same nodes and readout can be rebuilt on a two-dimensional array or a three-dimensional microwire montage by swapping the geometry the edges are drawn from (Fig. 1b).

**Figure 1:**
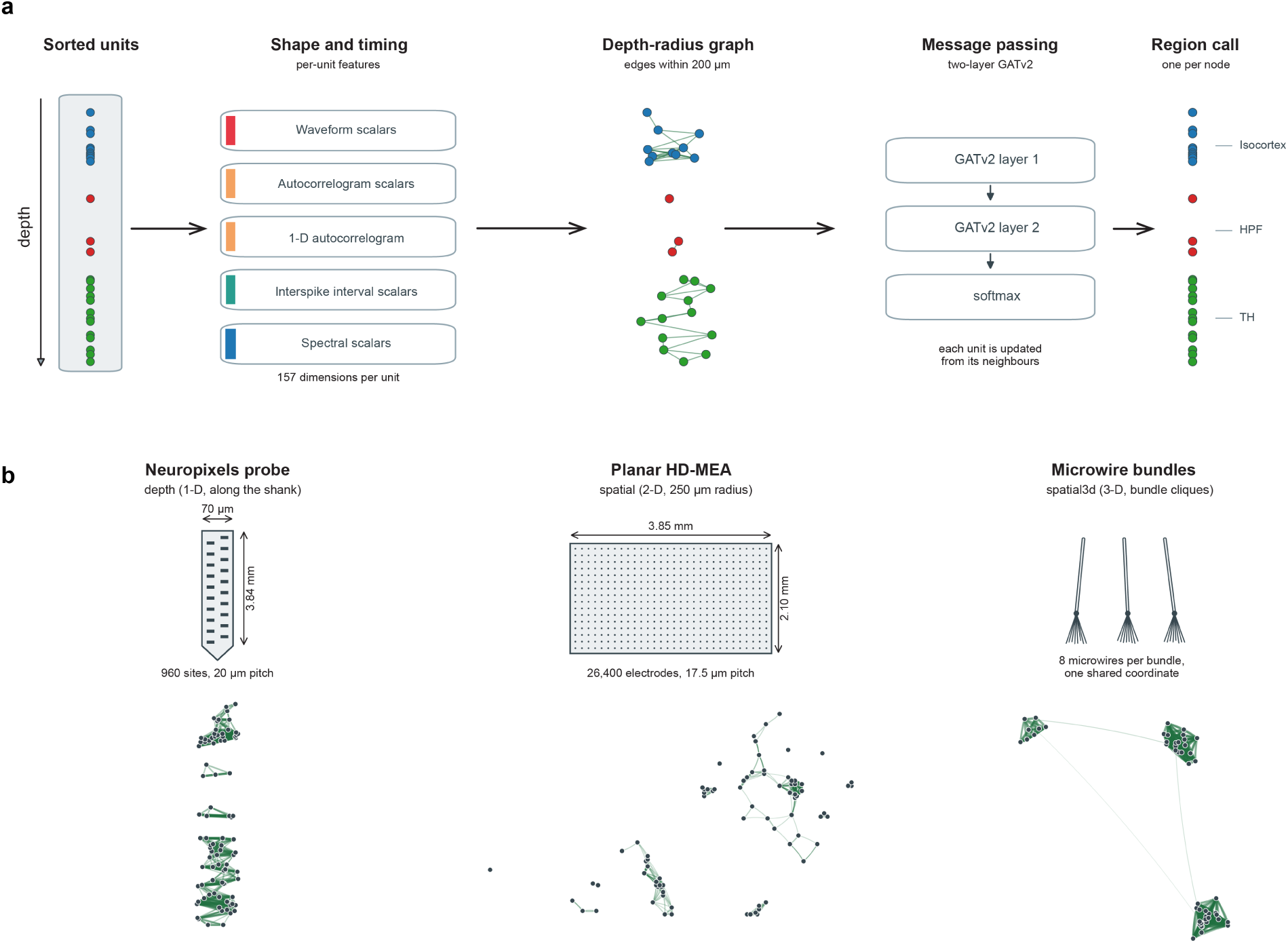
MYCEL composes per-unit features over a local population graph. **a**, Pipeline, drawn on one real IBL insertion. Sorted units on a probe each carry the same shape-and-timing features (57 waveform, interspike-interval, autocorrelogram and spectral scalars concatenated with the 100-bin autocorrelogram, 157 dimensions per unit); edges join units within a 200 *µ*m depth radius on the same probe; a two-layer GATv2 message-passes those features so each unit is updated from its neighbors; and a softmax returns one region call per unit. **b**, The same nodes and readout on three recording geometries, with only the edge relation rebuilt. These are depth along a Neuropixels shank, spatial within 250 *µ*m on a planar high-density multielectrode array, and spatial3d between microwire bundles, where each bundle shares one coordinate and so forms a clique. The shank’s lateral axis is exaggerated for display, and the microwires of each bundle are jittered apart.

### Local composition improves region decoding

We benchmark every method on matched units and leave-animal-out folds across five tasks. These are the Allen four-way super-region and its hippocampal-formation, thalamus and visual-area sub-region partitions^16^, and the IBL Brain Wide Map ten-way Cosmos partition^17^. The comparison spans the range of population context, from two single-unit encoders of intrinsic firing (metrics-LogReg and ISI-MLP)^6^ through a contrastively trained single-unit encoder (NEMO with *k*=1)^9^ to three methods that aggregate across units, NEMO with *k*=5 neighbor pooling, NuCLR^14^ and MYCEL (Methods). Unless stated otherwise every method is scored at a matched 90-min per-unit recording budget; the full method*×*task*×*budget grid is in Supplementary Fig. 6.

Single units carry regional signal, but it is modest and varies across tasks. On the Allen super-region (chance 0.25) the single-unit encoders reach 0.48 (metrics-LogReg) and 0.58 (ISI-MLP) balanced accuracy; on the finer sub-region tasks they fall toward chance, reaching 0.32–0.34 on the five-way hippocampal formation (chance 0.20), *∼*0.21 on the seven-way thalamus (chance 0.14) and 0.21–0.23 on the six-way visual areas (chance 0.17), while on the ten-way IBL Cosmos partition (chance 0.10) they clear chance roughly 2.5-fold (*∼*0.25). The single-unit signal is therefore strongest where the anatomical partition is coarse and weakest where neighboring areas are likely to share cell types (Supplementary Fig. 6).

Composing those features over the population improves on that baseline on every task (Fig. 2a). MYCEL has the highest mean balanced accuracy of any method benchmarked here, reaching 0.91 on the Allen super-region, 0.76 on the hippocampal formation, 0.41 on the seven-way thalamus, 0.49 on the visual areas and 0.53 on IBL Cosmos. These are 1.6 to 2.2 times the strongest single-unit encoder on each task. All four Allen super-regions are recovered (confusion matrices in Supplementary Figs. 7 and 8). Both published methods improve on the single-unit baselines on most tasks but fall short of MYCEL. The population model NuCLR reaches 0.58/0.68 (linear/*k*NN probe) on the super-region, 0.45/0.47 on the hippocampal formation, 0.18/0.26 on the thalamus and 0.25/0.33 on the visual areas. The contrastive encoder NEMO (*k*=5) is the closest competitor on the Allen partitions, reaching 0.87 on the coarse super-region, but the gap widens on the finer partitions (0.58 against 0.76 on the hippocampal formation, 0.33 against 0.49 on the visual areas). On identical leave-animal-out folds, MYCEL outperforms NEMO in all five folds on the super-region (*P* = 0.031), the hippocampal formation (*P* = 0.0006) and the visual areas (*P* = 0.012; two-sided paired *t*-tests, *n* = 5). On the thalamus the two are tied (0.41 against 0.40, *P* = 0.88).

**Figure 2:**
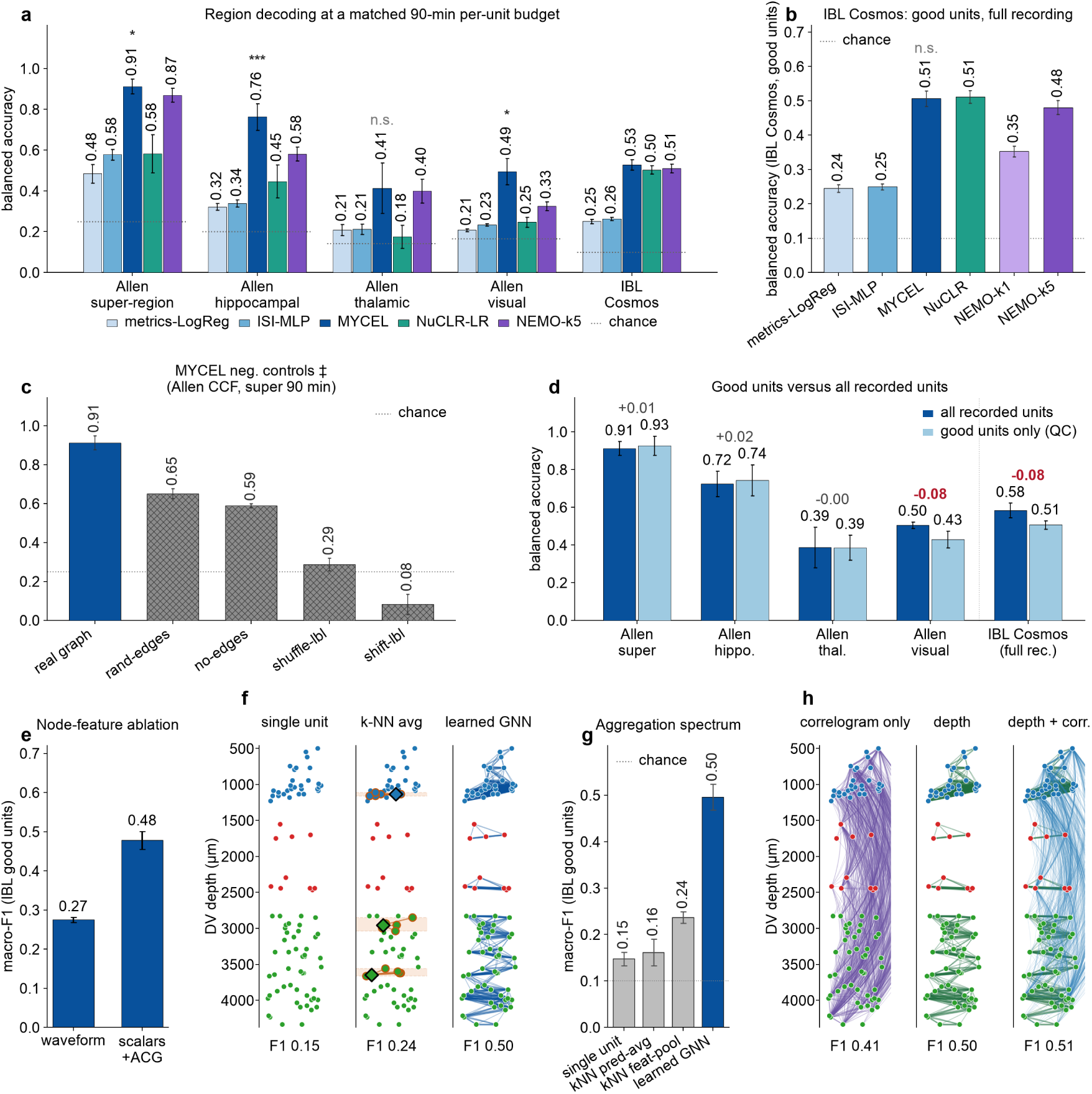
Region decoding performance, controls and sources of the gain. **a**, Balanced accuracy (leave-animal-out, inductive) on the Allen four-way super-region and its hippocampal formation, thalamus and visual areas partitions, and on the IBL ten-way Cosmos partition, every method at the same 90-min per-unit recording budget (IBL, good units from a sweep separate from **b**; NEMO, mean of the two folds completed at this budget). Dotted segments, per-task chance. Asterisks, MYCEL versus NEMO (*k*=5), two-sided paired *t*-test over the five folds (*^∗^P <* 0.05, *^∗∗∗^P <* 0.001; n.s., not significant). All budgets are in Supplementary Fig. 6. **b**, IBL Cosmos good units at full recording time. **c**, Negative controls on the Allen super-region at the same budget, showing the real graph, random edges, no edges, label-shuffle and label-shift. **d**, MYCEL on every recorded unit versus on the QC-passing “good” subset alone, for the four Allen tasks (super-region at 90 min, the others at 60 min) and for IBL Cosmos at its full recording time. The two arms score different unit sets (matched comparison in Supplementary Fig. 10). **e**, Node features, comparing bare waveform with the shape-and-timing features (mean *±* s.d., four initializations). **f**, Aggregation, drawn on one representative insertion (*n* = 88 units at true depths, colored by Cosmos region), showing single unit, *k*-NN depth average (*k*=5) and learned graph. **g**, That spectrum quantified. **h**, Edge definition on the same insertion, comparing correlogram-only, depth and depth+correlation. In **f** and **h** the macro-F1 beneath each panel is the method’s five-split mean on the matched IBL good units (Supplementary Table 1), not a score on that insertion. Center, mean; error bars, s.d. over folds.

Published population models^9,14^ report on the IBL corpus at full recording time, so we run the benchmark there as well (Fig. 2b). We re-ran MYCEL and NuCLR end to end on the identical 62,856 units passing quality control (QC) under five leave-animal-out splits. The two are statistically indistinguishable, tying at 0.51 balanced accuracy against 0.48 for NEMO (*k*=5) and *∼*0.25 for the single-unit encoders. The leaderboard macro-F1 metric agrees (0.476 *±* 0.055 against 0.453 *±* 0.013, two-sided paired *t*-test on the matched splits, mean paired difference +0.022, 95% CI *−*0.046 to 0.090, *t*_4_ = 0.90, *P* = 0.42, paired Cohen’s *d_z_*= 0.40, *n* = 5 splits; Methods). MYCEL reaches this parity with far less computation, because it combines each unit only with units within 200 *µ*m in depth, whereas NuCLR attends to every unit in the session. The cost comparison is in the Methods (Supplementary Fig. 9).

Negative controls show what each part of MYCEL contributes (Fig. 2c). On the Allen super-region at the same budget, destroying the region labels removes the signal. Label-shuffle falls to 0.29, near chance (0.25), and label-shift falls to 0.08. Label-shift falls below chance because the model learns the shifted labels. Removing the graph edges but keeping the node features gives 0.59, and randomizing the edges gives 0.65. Both are well above the label controls but well below the 0.91 of the real graph, so the node features and the graph each contribute.

Standard quality control discards roughly two-thirds of the Allen units and, by preferentially culling low-firing ones, can bias the surviving population toward particular cell types. Filtering to good units is close to neutral on the coarse Allen partitions (+0.014 super-region, +0.019 hippocampal formation) but costs 0.075 on the visual areas and 0.077 on IBL Cosmos (Fig. 2d). Because these two arms score different unit sets, we also ran a matched comparison. Returning the non-good units to the graph as context-only nodes, while scoring the *identical* good units, raises good-unit macro-F1 from 0.488 to 0.576, higher in every one of five splits (Δ = +0.088 *±* 0.064, 95% CI 0.008 to 0.168, paired *t*_4_ = 3.06, *P* = 0.038, Cohen’s *d_z_* = 1.37, *n* = 5; Supplementary Fig. 10). Repeating the full Allen benchmark on QC-filtered units shows that the decode does not depend on low-quality units. MYCEL still outperforms NuCLR and the single-unit encoders on all four tasks and the label controls fall to near chance (Supplementary Fig. 11).

We used three ablations to test whether the improvement comes from the node representation, its composition over the graph or the edge relation. First, we compared node features (Fig. 2e). On the matched good-units set the same architecture with only waveform nodes reaches macro-F1 0.275*±*0.006 against 0.478*±*0.023 for the shape-and-timing features on the identical split (four initializations of one split). Second, we compared aggregation methods (Fig. 2f,g). Over the identical depth neighborhood the learned graph reaches macro-F1 0.496 (a separate five-split sweep; Supplementary Table 1), against 0.24 for feature mean-pooling and 0.15 for the single unit. Averaging features across neighbors helps, but learned message passing does better, and no parameter-free aggregator matches it (Supplementary Fig. 12). Third, we compared edge relations (Fig. 2h) and found that most of the gain comes from depth edges. Co-firing-only and cross-correlogram-only graphs both fall below depth alone, and adding either relation on top of depth does not improve it substantially. We therefore use the depth graph and report functional edges only as a control. The choice does not depend on the graph architecture, as GCN^19^, GraphSAGE^20^ and a deeper GATv2 all land within one between-split standard deviation (Supplementary Table 1), and the decode is robust to the subgraph window, class weighting and most node-feature sets (Supplementary Fig. 13).

The decode also transfers between atlases. Trained on IBL and applied to Allen with no adaptation and no target labels in the shared Cosmos label space, MYCEL reaches 0.753 *±* 0.031 balanced accuracy on the four Cosmos classes both corpora sample in quantity (chance 0.25), and 0.516 *±* 0.042 across all six (*n* = 3 seeds; chance 0.167). The six-class figure is lower because Allen contributes only 12 cerebral-nuclei and 143 hypothalamic units, and these two classes are undecodable in either direction (recall *≤* 0.09). Because the two datasets differ in laboratory, rig, spike sorter, quality-control pipeline and behavioral paradigm, the transfer is unlikely to rely on dataset-specific activity statistics. The reverse direction is weaker on the same four classes (0.504 *±* 0.011), which we attribute to region coverage, since a model trained on Allen never sees the IBL regions that Allen does not record (Supplementary Fig. 14).

Each prediction also carries a confidence score that, after temperature scaling, is calibrated and tracks correctness. Summing the fine-level softmax over each super-region’s children returns a coarse call more accurate than the fine one (0.87 versus 0.63 on IBL Cosmos, 0.95 versus 0.45 on Allen), so when the fine call is uncertain, the unit can still be assigned at the coarser level (Methods; Supplementary Figs. 15 and 16).

### Embeddings recover anatomy without labels

MYCEL’s learned per-unit and per-insertion embeddings organize by Cosmos region (Supplementary Fig. 17). Unsupervised Leiden clustering of the same embeddings recovers this anatomy across the brain on IBL without using labels, as seen by placing each community back at its Allen Common Coordinate Framework (CCF) coordinates (Fig. 3). On the Allen Visual Coding corpus, the communities align only weakly with anatomy (Supplementary Fig. 18).

**Figure 3:**
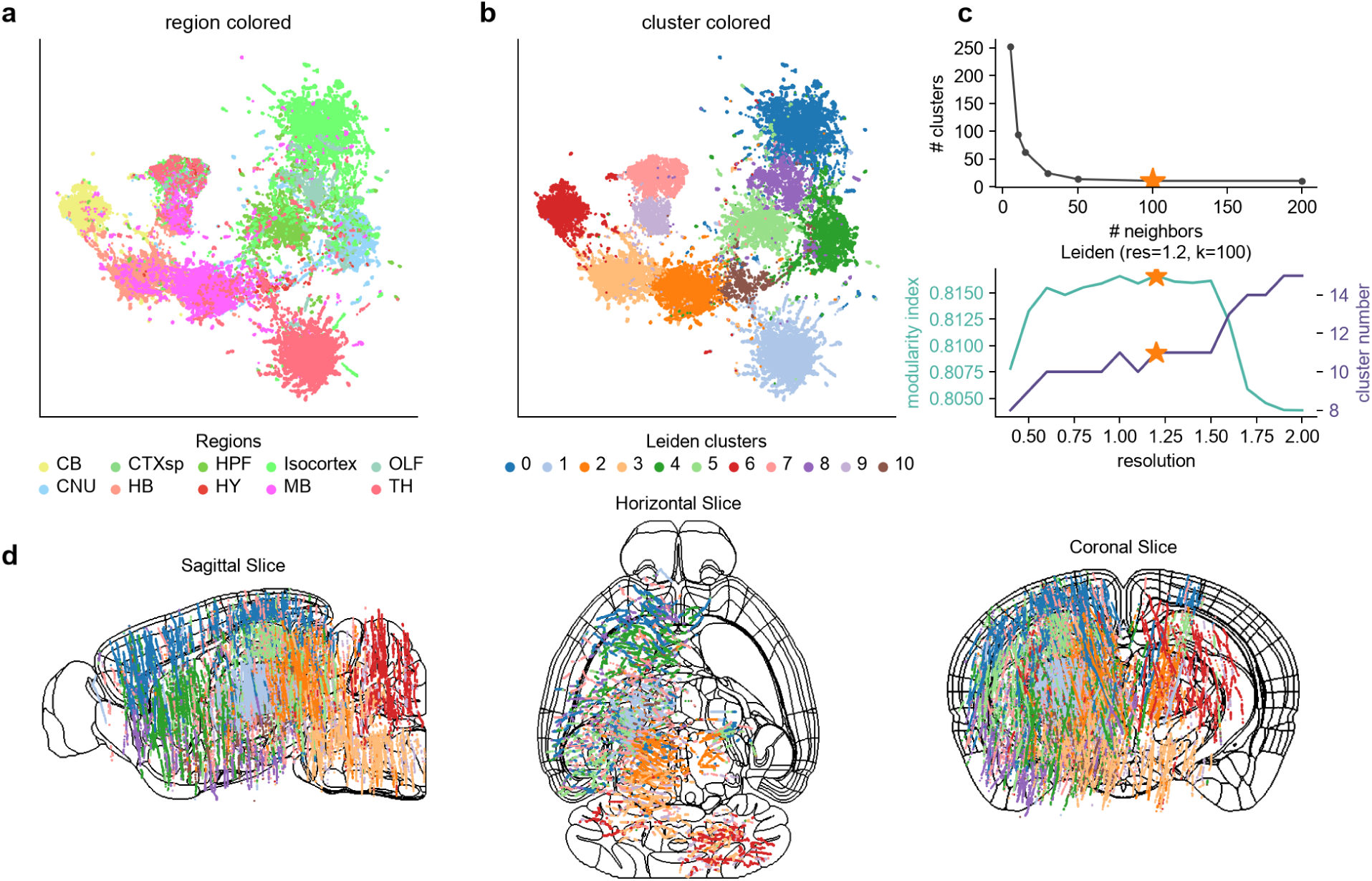
Unsupervised structure of the brain-wide IBL Cosmos embedding. **a**, Uniform manifold approximation and projection (UMAP) of the per-unit MYCEL node embeddings (*n* = 62,700 good units), colored by the 10-region Cosmos taxonomy. **b**, The identical UMAP geometry colored by *unsupervised* Leiden community of the embedding’s *k*-nearest-neighbor graph. **c**, Leiden operating-point selection, showing community count versus number of *k*-NN neighbors (top), and modularity index and community count versus resolution (bottom); stars mark the chosen point (neighbors = 100, resolution at maximum modularity), yielding 11 communities. **d**, Each unit placed at its Allen CCF coordinate on sagittal, horizontal and coronal boundary slices, colored by its Leiden community.

### Embeddings recover within-structure position

The embedding also carries continuous position within a structure. We tested this by predicting each unit’s Allen CCF coordinate from its embedding under a *leave-animal-out* split, so the regression cannot exploit probe or session batch (Fig. 4; controls in Supplementary Fig. 19). A sweep over every structure with enough units (Supplementary Fig. 20) reveals two reproducible within-structure gradients, each shared across anatomically related areas. The first is a cortical depth gradient along the dorso-ventral axis. The embedding recovers a unit’s depth on held-out animals in primary and anteromedial visual cortex (VISp *R*^2^ = 0.30, VISam 0.44), and the signal remains after subtracting each probe’s mean depth (within-penetration relative depth 0.34 and 0.42 against a shuffle-probe control near zero). It is absent in other visual areas (VISl, VISpm) and in the well-sampled thalamic nuclei (LGd, LP).

**Figure 4:**
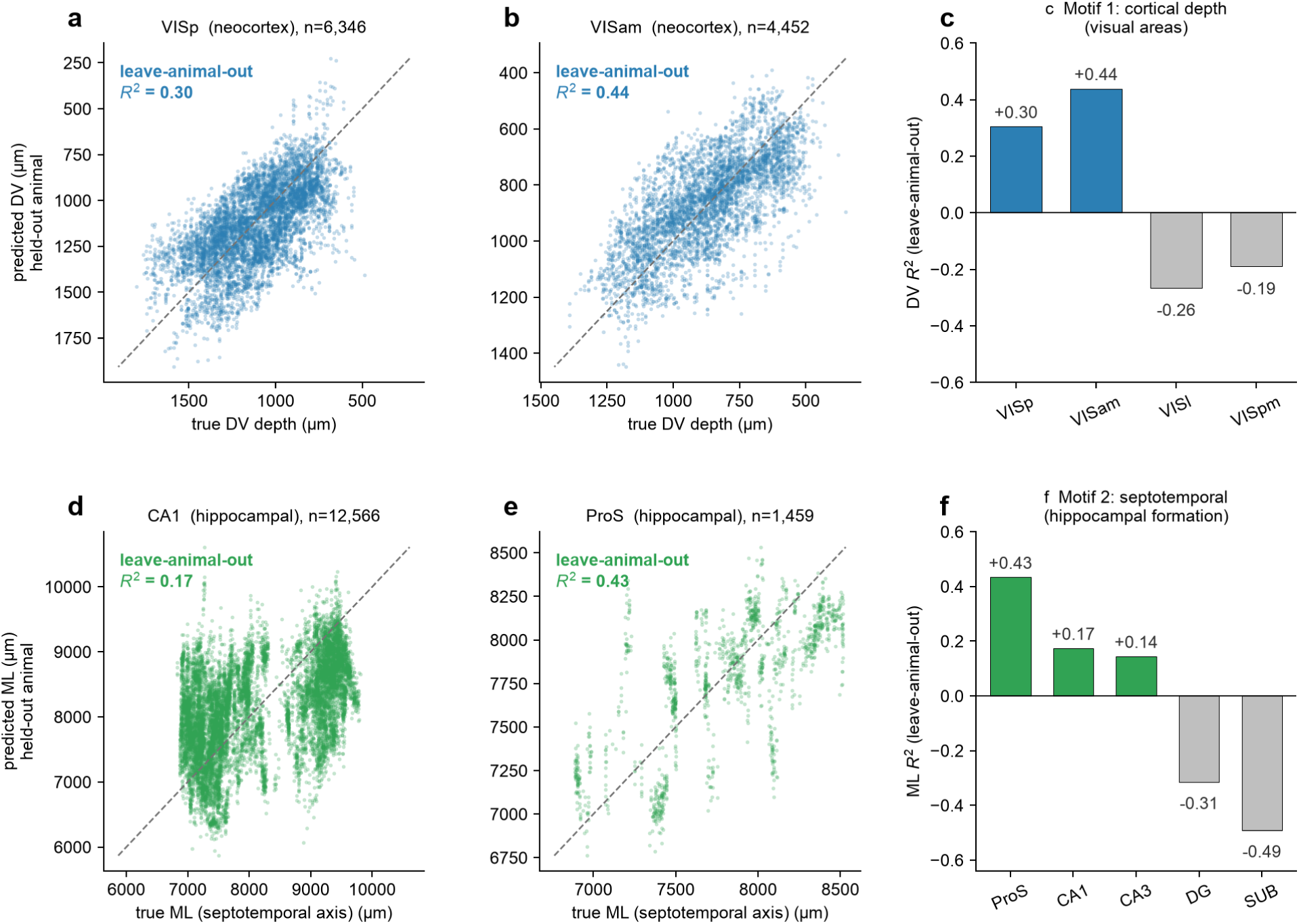
Two within-structure position motifs the embedding recovers on held-out animals. A ridge regression predicts each unit’s real Allen CCF coordinate from its 64-d MYCEL node embedding, with units grouped by the Allen-source structure_acronym (true anatomy, matched to the Allen source by exact unit identity). All *R*^2^ are out-of-fold under a *leave-animal-out* split of the regression (the embedding model was trained on 38 of the 47 animals; replication on the other nine in the main text), so the regression cannot exploit probe or session batch (controls in Supplementary Fig. 19; sweep over every structure with enough units in Supplementary Fig. 20). **Motif 1, cortical depth** (dorso-ventral, DV). **a**–**b**, predicted versus true DV depth (dorsal up) for VISp and VISam; **c**, DV *R*^2^ across visual areas. **Motif 2, hippocampal septotemporal position**, read out as the mediolateral (ML) coordinate. **d**–**e**, predicted versus true ML for CA1 and prosubiculum (ProS); **f**, ML *R*^2^ across the hippocampal formation. Dashed line, identity; *n* units per structure in the panel titles.

The second gradient is the hippocampal septotemporal axis. In the mouse this long axis runs largely mediolaterally, and the mediolateral coordinate loads 0.82 on the principal axis of CA1 position, which captures 91% of the variance. The embedding recovers a unit’s mediolateral position across the hippocampal-formation pyramidal fields and prosubiculum (prosubiculum *R*^2^ = 0.43, CA1 0.17, CA3 0.14) but not the dentate gyrus or subiculum, with the ProS and CA1 signals flat from leave-probe-out to leave-animal-out. Because the mediolateral axis is tangential to the penetration (most of its variance is between probes, with almost no within-penetration spread), this signal does not come from position along the shank. It is, however, almost entirely a between-probe quantity, and Allen probes enter at stereotyped positions, so it is partly set by which probe recorded the unit, a confound that leave-animal-out does not remove.

Although the regression holds animals out, the embeddings come from a single MYCEL trained on 38 of the 47 animals, so the network saw most of the held-out animals during training. Scored only on the nine animals the network never saw, within-penetration depth replicates (VISp *R*^2^ = 0.31, VISam 0.37, against 0.34 and 0.42 above) and absolute VISp depth holds at 0.18, so the cortical depth gradient does not depend on in-sample embeddings. The hippocampal signal is small on those nine animals (CA1 *R*^2^ = 0.03, against a shuffle control of *−*0.04), but this test has only 26 CA1 penetrations, and prosubiculum was recorded in only two of these animals. We therefore report the hippocampal gradient as early evidence. It is consistent with the known septotemporal variation in hippocampal electrophysiology and gene expression^21,22^, but these recordings cannot separate it from probe placement.

### Decoding generalizes across species and preparation

We next retrained the same approach on three new settings, two new species (macaque ventral stream, human medial temporal lobe) and a new preparation (planar organoid arrays), and asked in each case whether the method works beyond mouse Neuropixels and whether the population graph improves on the single unit (Fig. 5).

**Figure 5:**
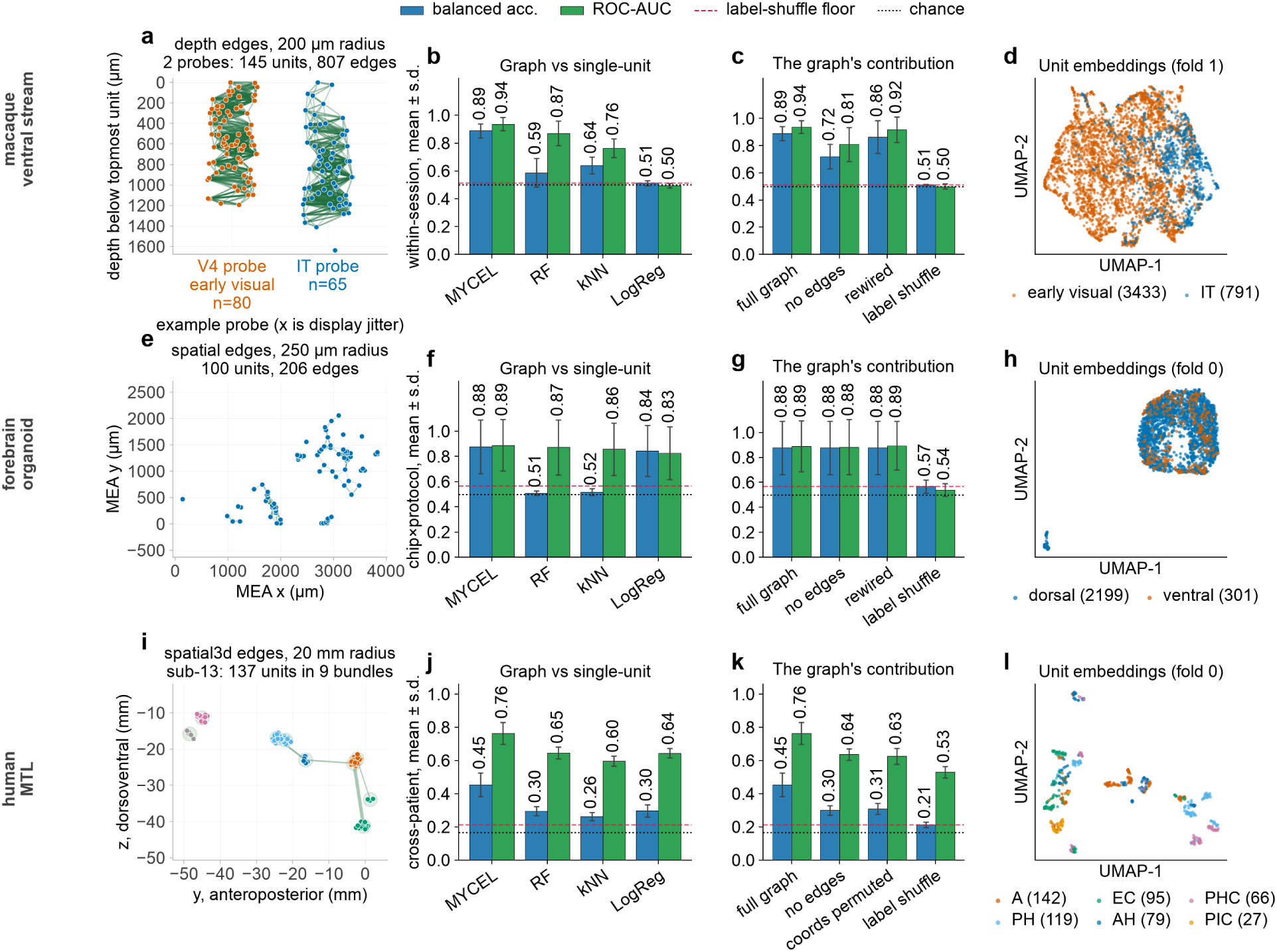
Decoding generalizes across species and preparation. The same shape-and-timing node features and population-graph readout, retrained on three new settings. Columns are parallel across rows and show an example graph; the decode against no-graph baselines, each row on the split that row supports (chip*×*protocol for the organoid, cross-patient for the human, and within-session for the macaque, whose cross-animal split has only two folds; Methods); graph contribution; per-unit embeddings. **a**–**d**, **Macaque ventral stream** (Triple-N ^23^; *n* = 19,130 units, 84 probes, 5 animals), IT versus early/intermediate visual cortex. **a**, Two single-area probes at cortical depth, depth-edge graph. **b**, Within-session decode against the matched-feature shape-and-timing baselines on identical folds, the same five-fold partition scored in **c**. The cross-animal split is in Supplementary Fig. 21a. **c**, Graph contribution (same within-session partition as **b**). **d**, Per-unit embeddings, held-out cross-animal fold. **e**–**h**, **Forebrain organoid** ( ^25^; *n* = 10,297 units, 94 recordings, *∼*21:1 imbalance), dorsal versus ventral patterning, panels as in **a**–**d** with the two-dimensional spatial graph and per-recording pooling. **i**–**l**, **Human medial temporal lobe** (SUMMER ^28^; *n* = 2,286 units, 29 patients), six-way region decode, three-dimensional spatial3d graph. A bundle’s microwires share one coordinate, so each bundle is a zero-distance clique (shaded in **i**) and only between-bundle edges are drawn. **k**’s randomized arm permutes which unit sits at which coordinate, rebuilding the graph at the identical edge count and degree sequence but reassigning which unit holds which degree; the macaque and organoid rows instead rewire the edge set at fixed per-unit degree (Methods). Center, mean; error bars, s.d. over folds. Dotted line, chance; dashed, the label-shuffle floor. The macaque cross-animal split, intrinsic timescales and four-way partition are in Supplementary Fig. 21.

On the Triple-N macaque ventral-stream resource^23^ we train a two-class MYCEL to call, from intrinsic electrophysiology alone, whether a unit is in inferotemporal cortex (IT) or early/intermediate visual cortex (EVC). The two classes separate well both within session and across animals (ROC-AUC 0.94 in both; Fig. 5a,b and Supplementary Fig. 21a). At the population level, autocorrelogram intrinsic timescales also run slower in IT than in EVC (median *τ* 62 versus 43 ms, two-sided Mann–Whitney *P* = 1.6 *×* 10*^−^*^163^), a shift between heavily overlapping distributions (per-unit ROC-AUC 0.67), extending the primate intrinsic-timescale hierarchy^24^ to the ventral stream. This difference is too small to explain the decode. Over the five-fold within-session partition MYCEL reaches balanced accuracy 0.89 and ROC-AUC 0.94, while the same shape-and-timing features read by single-unit classifiers reach at most 0.64 (Fig. 5b). Method comparisons on this dataset use that partition; the two-fold cross-animal split is reported descriptively (Methods; Supplementary Fig. 21a).

Most of this margin comes from the graph. Stripping every edge costs 0.17 balanced accuracy (0.89 *→* 0.72) against a label-shuffle floor of 0.51, and the finer four-way V1/V2/V4/IT partition depends on the edges more heavily still, falling from ROC-AUC 0.84 to 0.62 once depth edges are removed. Rewiring the edge set at fixed per-unit degree, which preserves every unit’s neighbor count while destroying locality, recovers most of the binary loss (0.86), so on this contrast having neighbors matters more than which neighbors they are (Fig. 5c). On a held-out cross-animal fold the per-unit embeddings separate the two classes (Fig. 5d).

The approach also transfers across recording preparation. Mouse forebrain organoids on planar MaxWell high-density multielectrode arrays (HD-MEAs)^25^ have no laminar axis to carry the depth edge, so substituting a two-dimensional spatial relation (units within 250 *µ*m on the electrode grid) is the only structural change (Fig. 5e). MYCEL then separates dorsal-from ventral-patterned organoids on four of five chip*×*protocol-grouped inductive folds (ROC-AUC 0.96–0.99) and is at chance on the fifth, which holds out only *∼*2 ventral recordings; the distribution is bimodal, so we quote folds rather than their mean (Fig. 5f). Dorsal (pallial) and ventral (subpallial) protocols yield sharply different excitatory and inhibitory censuses^26,27^. Here the graph does not help. Deleting all 107,582 spatial edges leaves the decode unchanged (0.877 with and without), as does rewiring them, while a label shuffle drops it to 0.567 (Fig. 5g). The per-unit features are already enough (Fig. 5f), and a class-balanced logistic probe on the same shape-and-timing features, with no graph, reaches balanced accuracy 0.845 against the full model’s 0.877. In the per-unit embedding the minority ventral units are interleaved with the dorsal ones (Fig. 5h). Because dorsal and ventral organoids also differ in cell line and acquisition batch, we read this as MYCEL transferring from Neuropixels shanks to a two-dimensional array, and not as a biological finding.

Finally, we applied the same approach to human recordings. In the SUMMER dataset ^28^ (DANDI:001616), 2,286 single and multi units were recorded on Behnke–Fried microwires in 29 intracranially implanted epilepsy patients, and we decode the six medial-temporal regions clearing a 50-unit floor across patients (Fig. 5i–l). Microwires are not a probe, so the graph is built from three-dimensional distance between recording sites; because a bundle’s eight microwires share one manually reviewed coordinate inside a single region, a distance graph groups same-region units by construction, so we tested any gain against structural controls. The graph improves decoding substantially, reaching balanced accuracy 0.455 against 0.297 for the strongest single-unit baseline, over a label-shuffle floor of 0.213 (chance 0.167; Fig. 5j), and against 0.300 for the identical node features with no edges (Fig. 5k). Permuting which unit sits at which coordinate, which rebuilds the graph at the same edge count and degree sequence but breaks the correspondence between a unit and its location, costs 0.146 and matches the no-edge arm, so the contribution depends on where units sit rather than on graph density (Fig. 5k). Deleting the within-bundle cliques outright still scores 0.391, so relations between bundles carry signal of their own. Edges built from co-firing rather than geometry add nothing (0.306). Pooling units by bundle, since each bundle lies in one region, reaches balanced accuracy 0.337 *±* 0.076 over 168 bundles, but held out by patient it falls to 0.202 *±* 0.029, indistinguishable from chance; we therefore report the bundle-level call as a within-patient result. Because a bundle’s own units do not identify its region across patients, the cross-patient signal in the unit-level decode comes from context between bundles. That context may be how physiology differs across the implant or the implant layout itself, which is similar from patient to patient; permuting coordinates removes both, so these data cannot separate them.

## Discussion

Extracellular spikes carry anatomical information at both the level of individual neurons and the local populations in which they are found. Single-unit electrophysiological features decode brain region above chance, consistent with previous evidence that spike timing contains anatomical signal^6^, but that signal becomes substantially stronger when the same features are interpreted across nearby units. Across the Allen partitions, local composition raises balanced accuracy by a factor of 1.6 to 2.2 over the strongest single-unit encoder on each task, and on the IBL Brain Wide Map it matches a dedicated population encoder under a matched protocol at a fraction of its cost. These features were chosen with a search on independent cell-type datasets and never tuned on brain region, so their transfer to anatomy is a finding in its own right. The electrophysiological signatures that distinguish cell types also carry information about location.

The gain from MYCEL does not come from extra model capacity, since parameter-free pooling over the same neighborhood stays below learned message passing, and scrambling the edges at matched capacity, edge count and degree sequence costs most of the gain. The likely explanation is the local distribution of neuronal types, which the graph reads alongside the type of the target neuron itself. Brain regions differ in cellular composition, laminar structure and circuit organization, so a population of electrophysiologically distinct neurons provides a richer anatomical signature than any unit by itself. This resembles spatial transcriptomics, where tissue domains emerge from molecular identities read together with their spatial context^29–31^, except that here identity is inferred from spiking. Testing it directly requires linking electrophysiological units to independent transcriptomic or morphological identities in the same tissue.

This hypothesis also predicts where the graph should help. Population context contributes most when the target distinction is poorly resolved by intrinsic electrophysiology, and little when it is already resolved. The two extremes are the mouse datasets (Allen and IBL), where the graph substantially improves classification, and the patterned forebrain organoids, where a single-unit classifier on the same node features nearly matches it and the classes stay separable with the graph removed entirely. How much the graph adds also depends on which single-unit representation it is measured against. Against hand-designed per-unit features the margin is largest where those features are weakest. AAgainst NEMO ^9^, whose contrastive per-unit embeddings are pooled over five neighbors and which is the closest competitor here, it holds on the super-region, hippocampal formation and visual areas partitions but closes on the thalamus, where the two are indistinguishable both on the unfiltered build (0.41 against 0.40, *P* = 0.88) and under quality control (0.39 against 0.40). A learned per-unit representation can therefore capture some of what local composition adds elsewhere.

The graph also keeps anatomical information at every spatial scale, from single units to the whole recording. This is important for Neuropixels data, because one insertion can cross several regions, and a single embedding per insertion would lose the boundaries between them.

The model also recovers anatomical information beyond discrete region labels. Within structures, the embedding preserves continuous position along cortical depth and along the hippocampal septotemporal axis, both axes on which neuronal electrophysiology and molecular identity are known to vary^21,22,32^. Cortical depth replicates on animals held out from both the network and the regression, whereas the hippocampal gradient remains early evidence because it cannot yet be separated from probe placement. The closest prior result resolves cortical depth categorically, assigning units to cortical layers from interspike intervals^6^; here we predict depth as a continuous coordinate on held-out animals. Because these predictions are Allen CCF coordinates, they can be compared directly with gene expression gradients registered to the same atlas, which could help test whether the gradients reflect neuronal identity, local circuit organization, physiological state, or some combination.

Population context further allows the model to use units that standard analysis discards. Returning units that fail quality control to the graph as context-only nodes improves decoding of the good units in every split. A unit need not be isolated well enough to support a reliable individual prediction in order to inform the classification of its neighbors. This is useful in practice for large-scale extracellular recordings, where quality-control thresholds and spike-sorting pipelines differ substantially across laboratories and determine which units are retained at all.

Per-region performance appears to be limited mainly by data coverage. Per-region recall increases with the number of units a region supplies (*r*(log *N*) = 0.96). Regions with many units, such as thalamus and isocortex, reach about 0.8, while regions sampled by few insertions fall below 0.15. Calibrated confidence reflects the same quantity, so restricting predictions to the most confident units raises accuracy substantially, while balanced accuracy increases less. The method can therefore already be used at the coarse level and with confidence thresholds, and accuracy on undersampled regions should improve as more insertions are recorded there.

Where a probe track has been reconstructed and registered, a regional call derived from spiking complements that registration rather than replacing it. The two rest largely on different evidence, the geometry of the electrode against the physiology of the units recorded on it, so a unit whose predicted region disagrees with its registered label marks a position worth re-examining, either the alignment at that depth or the unit itself. Registration also limits what we can measure. Our region labels are themselves the output of a registration, so registration error enters the training signal and part of what is scored as a wrong prediction is a wrong label; the accuracies reported here may be lower than the accuracy attainable against perfect anatomy. Because the two corpora were registered by separate laboratories under separate pipelines, the cross-atlas transfer suggests that region decoding does not depend on the alignment conventions of either.

The framework transfers from mouse Neuropixels to macaque cortex, planar organoid arrays and human medial temporal lobe microwires, with only the spatial relation used to build the graph changing between preparations. Each setting tests a different kind of transfer, so these results speak to the portability of the approach; they do not show that one biological mechanism underlies all of them. Where no atlas registration exists, as in organoid arrays, a calibrated regional call derived from spiking supplies an annotation that is otherwise unavailable; in human microwire recordings, localized from imaging and reviewed implant targets rather than a registered probe track, it provides an independent check.

Our approach is complementary to end-to-end population representation models such as NuCLR^14^, which learns neuron representations from population activity, whereas MYCEL begins with explicit electrophysiological features and composes them through sparse local message passing. That two substantially different representational strategies extract comparable information on the matched IBL benchmark, with the sparse and interpretable one scoring an order of magnitude fewer pairwise interactions, is consistent with the coverage result above, which suggests that both are limited more by the data than by their architecture. Representation learning for extracellular electrophysiology has otherwise focused on learning latent spaces from raw waveforms, derived single-unit features or population activity ^9,14,33,34^. Our feature search addresses the narrower question of whether the per-unit representation should itself be treated as fixed. Holding the evaluator constant while varying the input representation shows that useful feature combinations differ across cell-type datasets, although autocorrelogram-derived quantities recur across them and have been emphasized by independently designed methods^8,9^. We do not claim that the selected features are a universal electrophysiological code, only that features chosen without reference to anatomy still work well in a population graph. The two strategies are not exclusive. A frozen unsupervised latent trained on the same corpus did not improve on the hand-designed features as node input to the full model (macro-F1 0.44 against 0.49 on one inductive split; Methods). This likely reflects the size of this corpus, and we would not expect it to hold for learned representations in general. A contrastive encoder of the kind NuCLR uses, trained self-supervised on far more recordings than any single benchmark supplies, may learn a per-unit representation that hhand-designed features cannot match, and the coverage result above suggests it will. A natural next step is to use such an encoder’s per-unit embeddings as node features and combine them over the same sparse geometric graph. One summarizes the activity of a whole session and the other a 200 *µ*m neighborhood, so combining them would separate how each neuron is represented from how neighbors are combined, and we expect the combination to outperform either alone.

Several limitations apply. The feature search is restricted to a hand-curated primitive library and a single held-fixed cVAE evaluator, so another candidate space or evaluator could yield a different representation^35,36^, and the library excludes local field potentials, which carry anatomical information of their own^37^. The opto-tagged cell-type datasets do not permit a clean leave-animal-out analysis because cell-type labels are confounded with animal or genetic line, and the selected features may reflect both stable neuronal properties and network state. Cell-type composition cannot be measured directly in the large anatomical corpora, so the compositional account remains an interpretation rather than a demonstrated mechanism. The anatomical analyses also have limits. The strongest cross-animal evidence comes from the mouse datasets, the macaque leave-EVC-animal-out comparison has only two folds and is read descriptively, and the organoid experiment supports transfer across recording preparation rather than any developmental claim. In the human recordings anatomical geometry enters graph construction directly, and patient-held-out bundle-level classification remains at chance. In the mouse graphs, by contrast, edges use only each unit’s position on the probe, so no atlas information reaches the model. Finally, the method requires a meaningful spatial relation between units. Probe depth, planar-array coordinates and microwire locations supply one here, but recordings without spatial information cannot exploit this form of local composition.

Taken together, these results indicate a hierarchical organization of anatomical information in extracellular electrophysiology, expressed both categorically and, within structures, as continuous topography. Features that identify cell type also help identify location, because intrinsic electrophysiology and local population organization together constrain where a neuron is, although neither determines it alone.

## Methods

### Data processing pipeline

All datasets are ingested through a single pipeline that writes one Neurodata Without Borders (NWB) file per dataset^38^ with a uniform schema that includes a per-unit peak-channel mean waveform, the multi-channel mean waveform across the peak channel and its *±*10 neighbors, the spike-time list, an animal/session identifier, optional layer/depth metadata, and the cell-type or brain-region label. Recording-time budgets are applied to these files. Per-primitive extractors read this NWB once and emit each of the 16 feature primitives at canonical fixed shapes (waveforms resampled to 90 samples; ISI histograms at 60 linear bins over 0–60 ms; autocorrelograms at 120 bins of *±*60 ms at 1 ms, with a 10 *×* 100 two-dimensional variant), so every candidate sees inputs of identical shape and only the active subset varies.

### Datasets

Two conventions run through this paper. *Good* units are those passing the standard spike-sorting quality control of their corpus; *all* units are every sorted unit the recording yields. For Allen this is the three AllenSDK-default criteria (interspike-interval violation *<* 0.5, amplitude cutoff *<* 0.1, presence ratio *>* 0.9), which retain between a quarter and a half of units depending on the task; for IBL it is the Brain Wide Map’s own “good” label, the subset on which published population models report, which retains 62,856 of 643,596 raw units. Because the two corpora differ in which set is conventional, the Allen region analyses run on all units and the IBL head-to-head on good units; every figure states which, and the effect of the choice is quantified in both directions in Fig. 2d and Supplementary Fig. 10.

- **Hull** (mouse cerebellum): *n* = 103 opto-tagged units from 9 mice (13 sessions), 5 labeled classes, from the C4 database^8^; classes with fewer than 10 cells are excluded from training and evaluation, so the two Golgi cells are dropped and the analyzed set is *n* = 101 units over 4 classes (MFB, MLI, PkC_cs, PkC_ss), at chance 0.25.
- **Lisberger** (macaque cerebellum): *n* = 668 expert-labeled units, 5 classes, from the C4 database.
- **Buzsáki/CellExplorer** (mouse cortex/hippocampus): *n* = 374 opto-tagged and juxtacellular ground-truth cells across six labels^39^. The probe is restricted to four classes by dropping the pan-GABAergic VGAT marker (*n* = 13), which is a superset of the parvalbumin, somatostatin and vasoactive-intestinal-peptide classes rather than a distinct type, and the small axo-axonic class (*n* = 29). This leaves *n* = 332 units over four classes (parvalbumin 183, somatostatin 114, juxtacellular 23, vasoactive intestinal peptide 12), at chance 0.25. Juxtacellular names the recording method rather than a molecular type; we keep it as the dataset’s own ground-truth label.
- **Allen Visual Coding** ^4,16^: *n* = 120,087 units from 58 mice, one session each, across 332 probe insertions. All Allen numbers below come from this one corpus. The four region tasks are label subsets of it, the QC criteria filter it, and the area-restricted build used for the embedding figures is the same recordings narrowed to 19 areas spanning visual cortex, hippocampal formation, thalamus and midbrain (61,770 units, 47 mice, 225 insertions). For the main figure no unit-quality filtering is applied. Before the QC the four Allen tasks carry 104,642 (super-region), 46,403 (hippocampal formation), 12,261 (thalamus) and 45,260 (visual areas) units, the super-region total exceeding the sum of the three sub-partitions by its 647 midbrain units and the 71 postsubiculum units its hippocampal class carries but the five-field hippocampal formation task excludes. As a validation, applying the three AllenSDK-default quality criteria (ISI-violation *<* 0.5, amplitude cutoff *<* 0.1, presence ratio *>* 0.9) retains 35,240 (super-region), 11,643 (hippocampal formation), 5,756 (thalamus) and 17,608 (visual areas) units, a retention ranging from 25% on the hippocampal formation task to 47% on the thalamus, bringing the super-region set to *∼*750 per mouse, within the standard-QC range. Region evaluation groups on the specimen (animal) identifier. Region labels are assigned by registering each unit to the Allen Common Coordinate Framework (CCF).
- **IBL Brain Wide Map**^17^: units at IBL quality label “good”, mapped to the 10-region Cosmos taxonomy, under a leave-animal-out split (75/10/15% of animals to train/val/test, no animal in two sets). The IBL “good” QC label reduces the 643,596 raw Brain Wide Map units (590,104 with a fine region label) to the matched good-units corpus of 62,856 QC-passing units (62,700 labeled) across 693 insertions / 139 animals.
- **Forebrain organoids (dorsal/ventral)**: *n* = 10,300 MaxWell HD-MEA units (9,845 dorsal / 455 ventral) across 95 recordings from 27 chip*×*protocol groups, of which one recording carries too few units to build a graph, so 10,297 units across 94 recordings enter the analysis; from mouse forebrain organoids^25^ (DANDI:001374), labeled by patterning protocol. Units are braingeneers auto-curation outputs (Kilosort2 followed by a low-ISI auto-curation pass); we apply only sanity filters (finite waveform, non-empty spike train, per-unit spike cap) on top. Pharmacology-condition recordings are excluded. The planar array has no depth axis, so each unit’s peak-electrode (*x, y*) coordinate wires the spatial graph edge, and cross-validation is grouped on chip*×*protocol (stratified group *k*-fold, five label-stratified folds, split at the recording level) so no organoid appears in both train and test.
- **Triple-N** (macaque ventral stream)^23^: *n* = 19,130 Neuropixels units across 84 probe insertions from 5 macaques, spanning four areas (V1/V2/V4/IT). Each probe samples a single area, so the area label is a whole-probe property. The binary contrast used for the cross-species generalization pools V1/V2/V4 into one early/intermediate visual class (*n* = 4,777) against IT (*n* = 14,353); the four-way partition retains the original areas. Cross-animal evaluation is leave-EVC-animal-out, which yields only two folds, so cross-animal orderings are read as descriptive.
- **SUMMER** (human medial temporal lobe) ^28^ (DANDI:001616, version 0.260702.0824): *n* = 2,286 units (1,190 single unit, 1,096 multi-unit) on Behnke–Fried microwire bundles from 29 intracranially implanted epilepsy patients, one session each, labeled with the manually reviewed implant target. Region decoding uses the six targets clearing a 50-unit floor (amygdala 559, entorhinal cortex 493, anterior hippocampus 418, posterior hippocampus 412, parahippocampal cortex 293, piriform cortex 68); the remaining 43 units stay in the graph as context and are masked out of the loss. Three source conventions are converted on ingest. Spike times are stored in milliseconds, not seconds. Waveforms are aligned so that sample 19 is the largest-magnitude sample, which for 2,246 of the units is a peak rather than a trough, so those are sign-flipped to the trough-negative convention. Finally, the electrode table’s axis descriptions are unmodified defaults, the values being MNI RAS millimeters. One graph is built per patient and edges are Euclidean distance between recording sites in all three axes (spatial3d, 20 mm radius, 10 mm decay); five-fold cross-validation at the insertion level is already fully cross-patient, because one insertion is one patient. Because all eight microwires of a bundle share one coordinate and a bundle lies in a single region, distance edges group same-region units by construction, so we report two structural controls alongside the decode: deleting the within-bundle cliques, and permuting which unit sits at which coordinate (Fig. 5k). The permutation leaves the set of recording positions untouched and rebuilds the graph from it, so the edge count and the degree sequence are preserved exactly while each unit inherits a different unit’s degree and neighborhood, but a unit’s physiology no longer matches its recorded position. This differs from the rewiring control used on the macaque and organoid rows, which keeps every unit’s own degree and randomizes only which units are joined. A second build makes probe_id the bundle rather than the patient so each graph carries a single region and the graph-level readout is defined. It is scored under both splits and the two are reported separately, as a five-fold partition over the 168 bundles, which is cross-bundle but within-patient, and a patient-held-out split. Only the second is a generalization claim.

### The cVAE evaluator

Each candidate encoding compiles into a multi-modal cVAE whose backbone family and training procedure are fixed across candidate encodings; only the set of active per-modality encoders, the fusion network and the total latent width reconfigure. Every active primitive is encoded by a modality-specific network, each emitting a 2*z*-dimensional head later split into posterior mean and log-variance. One-dimensional inputs (waveform, ISI, autocorrelogram, fine spike train, cross-correlogram) are encoded by a 1-D ResNet-18 (base width 64); the two-dimensional autocorrelogram by a three-layer 2-D CNN (base width 32); the multi-channel waveform by a four-layer 2-D CNN; the four scalar blocks and the local-population descriptor by a three-layer MLP (hidden 128); and the five spike-train binnings by distinct deep encoders. Those five feed a 1-D ResNet for the 1 ms train, a NuCLR-style transformer (*d*_model_ = 128, 20 ms bins) for the 60 s train, dilated temporal-convolution networks for the CEBRA-style and long binnings, and a leaky integrate-and-fire spiking encoder for the 1 ms train. A fusion MLP concatenates the active modality heads with the conditioning embedding and maps it through Linear *→* BatchNorm *→* LeakyReLU *→* Linear to a *z*-dimensional code, from which two linear heads form a diagonal-Gaussian posterior *q_ϕ_*(*z | x, c*). Per-modality decoders (a shared fully-connected stage followed by the modality’s transpose network) reconstruct each active input under a mean-squared-error likelihood. The per-modality latent width *z*_dim_ is a searched parameter in [8, 48] and the total latent dimension is *z*_dim_ *× n*_active_, so model capacity co-varies with primitive count by construction. Following HIPPIE^10^, the production presets route the class label only into the decoder (the encoder never sees it), and at evaluation the class embedding is replaced by zeros, so all reported probes are label-conditioning-free; the recording-technology embedding is a per-dataset constant and is retained, so it cannot carry within-dataset cell-type information.

Every candidate encoding’s cVAE trains with AdamW (learning rate 10*^−^*^3^, weight decay 10*^−^*^5^), batch size 128, gradient-norm clipping at 1.0 and early-stopping patience 30, minimizing the *β*-VAE evidence lower bound (ELBO)

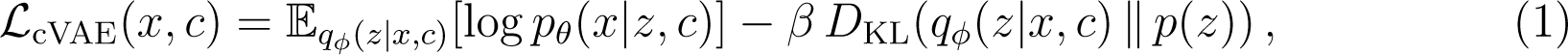

with a mean-squared reconstruction likelihood, a standard-normal prior and an analytic Gaussian Kullback–Leibler (KL) divergence *D*_KL_. The two search passes differ only in budget. Pass-1 trains each candidate’s cVAE for 60 unsupervised epochs on a single held-out fold; every non-penalty Pareto survivor is then retrained in pass-2 at the full HIPPIE-comparable budget (100 unsupervised plus 10 supervised epochs) under five-fold cross-validation. A categorical searched parameter selects among six conditioning and regularization presets that build on a plain *β*-VAE. These are a bare baseline (no conditioning, no fusion encoder, no normalization); the same with a class embedding and fusion encoder; a source-plus-decoder-only class-conditioning scheme; that scheme with batch normalization; the production default, which adds light input augmentation (probability 0.3, noise s.d. 0.03), class-embedding dropout 0.3, a reconstruction-consistency term and a five-epoch embedding warm-up; and a variant of the default with an auxiliary supervised contrastive loss on the latent mean.

### Evolutionary search

A candidate encoding carries 16 binary primitive flags, a categorical cVAE preset (six options) and two continuous parameters (*z*_dim_ *∈* [8, 48], *β ∈* [0.25, 4.0]). A non-dominated sorting genetic algorithm (NSGA-II)^35^ minimizes a four-vector consisting of the validation ELBO (*f*_1_); the negative balanced accuracy of a linear probe on the held-out fold (*f*_2_); the number of active primitives (*f*_3_); and log_10_ of a complexity proxy (*f*_4_). The search-time fitness probe is an L2-regularized logistic regression (*f*_2_); the headline cell-type numbers reported in figures instead use a cosine *k*-NN probe with *k* chosen on the training fold (matching HIPPIE’s native *k*-NN evaluation), while each comparison method is scored with its own published head (Classification probes, below). The complexity proxy *f*_4_ is an analytic, width- and latent-scaled estimate of model size, *f*_4_ = log_10_(8 *b*^2^*z n*_active_ + 4 (*z n*_active_)^2^ + 1) for backbone base width *b* and per-modality latent width *z*, not a measured floating-point-operation count; it orders candidates by capacity and is the quantity shown on the color axis of Supplementary Fig. 2b.

We run pymoo’s mixed-variable genetic algorithm (population 32, 16 generations, rank-and-crowding survival, random mating, duplicate elimination) in two passes. Pass-1 uses a custom comprehensive-sampling initializer in place of the i.i.d. sampler so that generation 1 covers every singleton primitive, every preset, and a curated set of published baseline encodings including HIPPIE’s waveform+ISI+autocorrelogram combination; it evaluates each candidate on a single held-out fold and writes the full non-dominated Pareto front. Pass-2 takes every Pareto survivor with non-penalty fitness (validation ELBO *<* 10^5^), retrains each at the full budget across all five cross-validation folds, and reports the swept-*k* cosine-*k*-NN balanced accuracy; pass-2 performs no further mutation, crossover or selection.

The search is the method’s one-time upstream cost, and we report it in Supplementary Table 2. Summing the recorded per-evaluation wall time over the production run, the three cell-type searches cost 130.5 h of aggregate evaluation time. Pass-1 is 512 candidate evaluations per dataset (20.4 h on Hull, 30.4 h on Lisberger, 28.3 h on Buzsáki; 79.1 h combined), and pass-2 retrains the Pareto survivors at full budget under five-fold cross-validation (115, 160 and 135 retrains; 0.7, 13.2 and 37.5 h; 51.4 h combined). Candidates were evaluated up to 32 at a time, so pass-1 completed in 7.8 h of wall clock. This cost is incurred once. The search is not repeated for any later experiment and is not included in the per-fold costs quoted for region decoding (Supplementary Fig. 9). The primitive library is in Supplementary Table 3; resolved operator settings are in Supplementary Table 4.

Seeding determines what the search can reach. On Buzsáki, a run initialized only with singleton and saturated encodings never evaluated HIPPIE’s three-of-sixteen combination within 16 generations of crossover, and so could not dominate it. This is why generation 1 is seeded with published baseline encodings. The optimal encoding also does not transfer across datasets. Hull is best served by scalar summaries alone (0.93), the larger Lisberger by HIPPIE’s trimodal histograms plus an autocorrelogram-scalar block (0.78), and Buzsáki by a waveform and two-dimensional-autocorrelogram combination distinct from both (0.63). No two datasets share a top-ranked candidate, and no transferable top candidate uses a population-context primitive (Supplementary Fig. 4). A latent-geometry counter-factual on the Hull and Lisberger search-selected cVAEs, and on a Buzsáki cVAE that includes the population primitive, confirms the composition (Supplementary Fig. 22). The autocorrelogram-derived scalar block is nevertheless selected on all three datasets. The node representation used in every later experiment is therefore a hand-assembled set built around that block rather than any single search output. The shape-and-timing features add the interspike-interval and spectral scalar blocks and a *±*100 ms, 2 ms-bin autocorrelogram histogram (Graph extension, below).

### Local-population-context primitives and leak controls

Two of the 16 primitives summarize a unit’s local population rather than its own spikes. The local_pop_stats primitive (10 features) collects, over each unit’s neighbor set, the mean and spread of pairwise 1 ms-binned Pearson correlation and of the spike-time tiling coefficient, a local participation ratio, the local firing-rate statistics, a network-burst coactivation fraction and the neighborhood size; the ccg_knn primitive (51 features) is the mean cross-correlogram (*±*25 ms at 1 ms) of the unit against the same neighbors. Both read a shared, session-cached neighbor computation. When a probe-geometry sidecar (per-unit probe identity and vertical position) is present, neighbors are the five nearest same-probe units within a 60 *µ*m vertical radius. When it is absent, as on the Buzsáki dataset, the neighbor set degenerates to five randomly chosen units from the same recording, and the swept-*k* probe then collapses to *k*=1, i.e. a within-session lookup.

Because every Buzsáki session is class-pure (one opto-tagged line per recording), this same-session neighborhood leaks the label through recording identity. We dissect it three ways. (i) A session-held-out split (group *k*-fold on session, every class still represented in train) drops local_pop_stats to balanced accuracy 0.48, at or below the per-unit-scalar reference. (ii) A neighbor-provenance ablation that draws the neighbor pool from *other* sessions collapses it to 0.30 (near the four-class chance of 0.25), while restricting to same-class other-session neighbors recovers only partially. (iii) The cross-validated probe selects *k*=1 on four of five folds for local_pop_stats but a larger *k* for the per-unit scalars, consistent with a same-recording lookup. We therefore plot the Buzsáki population descriptor only as a negative control and carry no population-context primitive into any transferable cell-type candidate.

### Graph extension (MYCEL)

Each sorted unit is a node. The headline node representation, the shape-and-timing features, is the 157-dimensional concatenation of the four hand-engineered scalar blocks (waveform 21, ISI 16, autocorrelogram 10, spectral 10; 57 total) with a 100-bin one-dimensional auto-correlogram histogram (*±*100 ms at 2 ms), z-scored per insertion when an insertion has at least 50 units; unit depth is deliberately *excluded* from node features to prevent the network from memorizing probe trajectories. Each node feature is projected to 64 dimensions by a small modality MLP. Two ablation node sets are compared against it: a bare 50-sample peak-channel waveform (the graph builds resample to 50 samples, not the 90 used by the search extractors) passed through a 1-D CNN, and a frozen unsupervised VAE latent (dimension 32) trained leakage-safe on the training-fold animals over the same 157-dimensional block. On the first IBL inductive split the VAE-latent nodes reach macro-F1 0.44 against 0.49 for the shape-and-timing features in an otherwise identical configuration.

Edges connect units within the same recording under one or more of four relations. These are depth (vertical separation *≤* 200 *µ*m, weight exp(*−|*Δ*d|/*200 *µ*m)); correlation (Pearson correlation of 20 ms-binned spike trains over a 60 s context *>* 0.05, weight equal to the correlation); xcorr (normalized cross-correlogram peak within *±*5 bins, i.e. *±*100 ms, exceeding 0.1, weight equal to the peak); and, for planar HD-MEA recordings, a two-dimensional spatial relation (electrode separation *≤* 250 *µ*m, weight exp(*−d/*150 *µ*m)). Edges are symmetrized and self-loops added. Depth edges are read directly from probe geometry, whereas both functional relations require binning every spike train and evaluating all pairs within a recording. These functional edges, as defined here, add nothing over depth; other formulations, such as a rate-corrected cross-correlogram, remain untested.

A two-layer GATv2^18^ (4 heads, hidden dimension 64, dropout 0.1; input projection and per-layer LayerNorm with residual connections) computes per-node region logits. Training is AdamW (learning rate 10*^−^*^3^, weight decay 10*^−^*^4^, gradient-norm clip 5.0) with cosine decay to 10*^−^*^5^ over 30 epochs (50 for the Allen tasks and the recording-duration sweeps), and the checkpoint is the epoch with the highest validation balanced accuracy. On the Allen and IBL leave-animal-out splits the validation animals are separate from the test animals, and the reported score is on the test animals. The macaque, organoid and human evaluations have no separate test set, so their reported score is the best epoch on the held-out fold itself, which is optimistic. Their label-shuffle arms are selected the same way and are the floor to compare against (dashed lines in Fig. 5); the single-unit baselines on those tasks involve no epoch selection. Splits are inductive at the animal level for the Allen and IBL region tasks and at the chip*×*protocol level for the organoids. The macaque within-session arm is a probe-level five-fold partition and is reported as such; its cross-animal counterpart is leave-EVC-animal-out. The default loss is unweighted cross-entropy. Inverse-frequency weights capped at 10*×* are used where the largest-to-smallest class ratio is severe, namely the hippocampal formation (8:1) and thalamus (8:1) sub-partitions and the organoid task (*∼*21:1), and not where it is mild, as in the six visual areas (2:1). The IBL Cosmos models are reported unweighted for comparability with the published baselines; Supplementary Table 5 gives the class-weighted alternative on IBL Cosmos, where weighting redistributes recall toward rare regions. For tasks where each probe or recording carries a single label (the single-area macaque probes and the organoid recordings) a graph-level readout concatenates mean and max pooling over the node embeddings and passes the result through a two-layer MLP head, yielding one prediction per graph; for node-level scoring that single prediction is broadcast back to its nodes.

The cost difference between MYCEL and the population model NuCLR follows from the two architectures. MYCEL is a two-layer GATv2 with 64 hidden units and four heads, 81,230 parameters in total, and it composes each unit only with the units inside a 200 *µ*m depth radius on its own probe. Its cost is therefore *O*(*nk*) in the *n* units of an insertion and the mean neighborhood size *k*, and the radius holds *k* at 11.2 edges per unit across this corpus (693 insertions, 62,856 units, 706,826 edges) independently of *n*. NuCLR attends all-to-all within a session, which is *O*(*n*^2^) in time and in attention memory. Over the same corpus that is 8,091,792 unit pairs against 706,826 edges, an 11.4-fold difference in the interactions scored, and the gap widens with insertion size, reaching sevenfold on the median 80-unit insertion and 35-fold on the largest at 391 units (Supplementary Fig. 9c). Measured per-fold wall times are in Supplementary Fig. 9a,b.

The single-unit baselines in the organoid and macaque method comparisons (Fig. 5b,f) are the matched-feature family, which use no population graph, the same node features MYCEL reads and the identical folds, so the gap to the GNN measures what message passing adds at matched features and matched held-out units. The metrics-LogReg and ISI-MLP encoders are the single-unit floor for the Allen and IBL region benchmark (Fig. 2a,b) and, of the cross-preparation tasks, are reported only on the macaque four-way partition (Supplementary Fig. 6). The matched-feature baselines apply three class-balanced classifiers to the same 157-dimensional shape-and-timing features. These are *L*_2_-regularized logistic regression (class_weight=balanced on standardized features), a random forest (300 trees, balanced_subsample) and a *k*-nearest-neighbor classifier (*k* = 15 on standardized features). On the organoid the split is the chip*×*protocol-grouped five-fold one and the class balancing addresses the *∼*21:1 dorsal:ventral skew. On the macaque the two splits are replayed from MYCEL runs themselves so the held-out sets are identical unit for unit. Within-session is the probe-level five-fold partition (a seeded permutation over the 84 insertions), and cross-animal is the leave-EVC-animal-out fold map (M1, M4); the recovered held-out unit counts match those recorded by MYCEL runs exactly. The organoid MYCEL itself is trained with an inverse-frequency class-weighted cross-entropy.

### Cross-atlas transfer

For the cross-corpus test (Supplementary Fig. 14) the model is trained on one corpus and evaluated on the other with its weights frozen, so no fine-tuning, target labels or target-corpus statistics enter the model. Both corpora are registered to the Allen CCF and coarsened to the ten-region Cosmos taxonomy, and scoring is restricted to the six classes both corpora contain (cerebral nuclei CNU, hippocampal formation HPF, hypothalamus HY, isocortex, midbrain MB and thalamus TH), so the comparison is over one label space. Scoring covers the 120,044 of 120,087 Allen units that carry a Cosmos-mappable label, the remaining 43 being ventricular. Predictions the source head can emit but the target corpus does not contain are retained and reported as an explicit off-taxonomy fraction rather than discarded. Node features are the shape-and-timing features and edges are depth, exactly as for the within-corpus region tasks; the recording-technology conditioning embedding is not used, so nothing in the model identifies the source corpus. Each direction is run at three seeds on the CCF-registered build at the 60-min per-unit budget, and the within-source ceiling quoted alongside each transfer is that same model scored on held-out animals of its own corpus in the same six-class space, which is not comparable to the four-way and ten-way values in Fig. 2.

### Confidence scores, calibration and hierarchical prediction

Every MYCEL prediction carries a confidence score, defined as the maximum class probability of the temperature-scaled softmax over the model’s output logits, *p*^ = max*_c_* softmax(*z/T*)*_c_*. The scalar temperature *T* is fit by minimizing the negative log-likelihood on an animal-disjoint validation split (never the test split), the standard post-hoc calibration of Guo et al. ^40^. We quantify calibration on the held-out test split with the expected calibration error (ECE), the sample-weighted mean gap between confidence and accuracy across 15 equal-width confidence bins, reported before and after temperature scaling, alongside reliability diagrams (Supplementary Fig. 15). Because the region taxonomies are hierarchical, the same fine-level softmax yields a prediction at any coarser anatomical level by summing the probabilities of that node’s children. For IBL we group the ten Cosmos regions into the three Allen top-level divisions (cerebrum, brainstem, cerebellum) and for Allen the 19 areas into the four super-regions (visual, hippocampal, thalamic, midbrain). The coarse prediction inherits a calibrated confidence the same way, and we report the fraction of fine-level errors that nonetheless fall in the correct super-region (“graceful” errors) as a hierarchy-aware error measure. Temperature scaling and hierarchical aggregation are pure post-processing of the trained model’s logits; they change neither the model nor its hard predictions.

On IBL Cosmos the raw logits are over-confident and temperature scaling cuts the expected calibration error from 0.166 to 0.025; on Allen they are already well calibrated (*T* = 1.14, calibration error 0.049). Confidence separates correct from incorrect calls (IBL Cosmos mean 0.71 correct versus 0.50 incorrect), and 68% (IBL) and 93% (Allen) of fine-level errors are graceful. Selective prediction is therefore useful, since retaining the most-confident 25% of IBL units raises accuracy from 0.63 to 0.91, plateauing near 0.93 on the top 10% and 5%, with per-region confidence tracking per-region recall (Spearman *ρ* = 0.67). The gain is larger in plain than in balanced accuracy (the latter peaking near 0.66) because confidence correlates with how many units a region supplies, so thresholding favors well-sampled regions.

### Intrinsic timescale

The per-unit intrinsic timescale *τ* (Supplementary Fig. 21b) is the decay constant of a single-exponential fit *A* exp(*−t/τ*) + *B* to the variance-normalized spike-count autocorrelation, computed in 50 ms bins over lags of 50–1000 ms; lag 0 is excluded to avoid the refractory/adaptation dip, following Murray et al.^24^. A unit contributes a *τ* value only if it has at least 50 spikes, the recording yields at least 40 bins, and the exponential fit reaches *R*^2^ *≥* 0.5, leaving *n* = 14,600 fit units. The *τ* distribution is shifted slower in IT than in early/intermediate visual units (median 62 vs 43 ms; IT-versus-EVC ROC-AUC 0.67; two-sided Mann–Whitney *U* test, *U* = 2.17 *×* 10^7^, *P* = 1.6 *×* 10*^−^*^163^, *n*_IT_ = 11,858, *n*_EVC_ = 2,742).

### Classification probes

Each method is evaluated with the probing head used in its own published benchmark, so that every bar reflects that method’s native evaluation. Our *k*-NN probe uses cosine distance on L2-normalized embeddings with *k* selected by cross-validation *on the training fold* only, then applied once to the held-out fold; HIPPIE is evaluated with its native *k*-NN (cosine distance, as in its release), NuCLR with a linear and a five-fold *k*-NN probe on its embedding, NEMO with its multilayer-perceptron probe, PhysMAP with its K-NN, and LOLCAT with its end-to-end set-attention classifier. Where we run standard probes ourselves the linear probe is unweighted logistic regression and the MLP probe uses hidden sizes (256, 128) with early stopping. The IBL Cosmos panel compares our end-to-end MYCEL against the population model NuCLR, the single-unit contrastive encoder NEMO, and two single-unit intrinsic-firing baselines (metrics-LogReg and ISI-MLP).

### Statistics and reproducibility

We report balanced accuracy as the primary metric and macro-F1 as secondary, and quote the other metrics alongside whenever a result is headlined. Two tasks also report a second metric for comparability with prior work. The IBL Cosmos region task reports macro-F1, the metric on which the NuCLR and NEMO leaderboards we benchmark against are reported; and the cross-preparation tasks also report ROC-AUC, which does not depend on a decision threshold (Fig. 5). Balanced accuracy is the mean per-class recall; macro-F1 is the unweighted mean of per-class F1. We use two split protocols. Cell-type evaluation is *transductive*, using stratified five-fold cross-validation at the unit level (seed 42), matching the HIPPIE benchmark, so units from one recording can fall in both the training and test folds. Brain-region evaluation is *inductive*, and every fold holds out whole animals (Allen and IBL) or whole recordings (organoids), so the model is tested only on animals or recordings absent from training. The one exception is the macaque within-session arm, which is a probe-level five-fold partition; its cross-animal counterpart (leave-EVC-animal-out) is animal-disjoint like the rest. Animal-level splits are needed because units within an animal share either a probe or the neural context, and a unit-level split would let that shared context leak from training into test. The IBL inductive split is the animal-disjoint protocol, audited for train/test overlap before training; the Allen split is grouped on the specimen identifier and the organoid split on chip*×*protocol. Significance tests are two-sided and are reported, with effect sizes, only where a *P* value is quoted in the text. These are Welch’s *t* for the between-method cell-type comparisons on Lisberger and Hull (Supplementary Fig. 2d); a paired *t* for the IBL Cosmos macro-F1 head-to-head against NuCLR (both models re-run on the identical five leave-animal-out splits) and for the context-only ablation; a paired *t* for the four Allen head-to-head comparisons against NEMO (*k*=5), the closest competitor on those tasks, computed over the identical five leave-animal-out folds at the 90-min budget (super-region, mean paired difference +0.043, 95% confidence interval (CI) 0.006 to 0.079, *t*_4_ = 3.26, *P* = 0.031, Cohen’s *d_z_*= 1.46, higher in 5 of 5 folds; hippocampal formation, +0.182, CI 0.131 to 0.233, *t*_4_ = 9.90, *P* = 0.0006, *d_z_* = 4.43, 5 of 5; visual areas, +0.169, CI 0.062 to 0.276, *t*_4_ = 4.38, *P* = 0.012, *d_z_* = 1.96, 5 of 5; thalamus, +0.014, CI *−*0.213 to 0.240, *t*_4_ = 0.17, *P* = 0.88, *d_z_* = 0.07, 2 of 5, which we read as a tie); and a Mann–Whitney *U* test for the macaque intrinsic-timescale contrast. These are pre-specified head-to-head comparisons against a named competitor on partitions fixed in advance, so no multiple-comparison correction is applied and none of them is used to establish a result on its own. Elsewhere, we did not run formal significance tests, and comparative words (“higher”, “lower”, “strongest”) denote differences in fold-mean balanced accuracy reported as mean *±* s.d.; we call two methods comparable when their means fall within one between-fold s.d. and separated when they do not, and given the small sample sizes (typically *n* = 5 folds, *n* = 2 for the macaque cross-animal split) all such differences are descriptive rather than inferential. Sample sizes (*n*) denote folds, repeat runs, or inductive splits as stated per figure. Several configurations were run as more than one independent five-split sweep. Each figure reads a single sweep throughout, so a method bar and the controls beside it always come from the same sweep, and values are compared within a figure rather than across figures; where a panel reads a different sweep from the one its neighbors read, the legend says so.

### Methods benchmarked against

Every published method is run from its own released implementation on the identical unit sets and folds, at the hyperparameters of its release. The two single-unit intrinsic-firing baselines are our own reimplementations of the published method. For MYCEL no hyperparameter search was performed. The configuration used in every benchmark comparison is the first one tried, and the graph-backbone and network-size sweeps were run post hoc as controls rather than to select it. Three settings vary by task (epoch budget, class weighting and the checkpoint reported), and are stated where they apply. The methods, and what each one consumes, are as follows.

- **HIPPIE**^10^: cVAE fusing the mean waveform, the ISI distribution and the autocorrelogram into a single per-unit embedding, run at the locked hyperparameters of the HIPPIE benchmarking release and evaluated with its native *k*-NN probe.
- **NEMO**^9^: contrastive single-unit encoder aligning the peak-channel waveform with a two-dimensional autocorrelogram, run at the release hyperparameters, with its native MLP probe on cell type and *k*-NN probes (*k* = 1, *k* = 5) on region.
- **Baseline cVAE**: the bare *β*-VAE preset of the same encoder family used by the search (no conditioning, no fusion encoder, no normalization), run at the HIPPIE-release hyperparameters and probed with the common swept-*k* cosine-*k*-NN.
- **PhysMAP**^7^: weighted-nearest-neighbor multi-view embedding over the mean waveform, ISI distribution and derived scalars, run at its release settings with its native *k*-NN probe.
- **Waveform random forest (WF-RF)**^41^: random forest on hand-engineered mean-waveform scalars, the fixed-encoding waveform reference, run at the HIPPIE-release settings.
- **LOLCAT**^11^: spike-train-only inter-spike-interval attention classifier, run from its released implementation. Each unit’s spike train is divided into 3 s snippets, each snippet reduced to a log-inter-spike-interval histogram, and the resulting set pooled by the model’s multi-head attention into a per-unit prediction (cell type only, five-fold). It carries no waveform or population input, making it a dedicated single-unit spike-timing reference for the cell-type task. The authors specify that snippets may be collected from contiguous recordings where trials are absent. We verified that none of the three cell-type datasets carries trial structure. The C4 export exposes no trial, stimulus or event field for Hull or Lisberger, and across all 86 CellExplorer ground-truth sessions 63 carry no epoch table and the remaining 23 carry a single epoch spanning the whole recording, with no session carrying more than one. The only stimulus-locked fields in that corpus are the optogenetic tagging pulses, which define the ground-truth labels and so cannot be used to align a classifier without leaking them.
- **metrics-LogReg**: single-unit intrinsic-firing baseline, our reimplementation of the features and readout used to report single-unit anatomical codes ^6^. Fourteen scalar spike-train metrics are computed per unit from spike times alone. These are log firing rate, log inverse first-percentile ISI, log median ISI, log mean ISI, ISI coefficient of variation, CV_2_, local variation LV, burst fraction (fraction of ISIs *<* 6 ms), Fano factor of 100 ms bin counts, presence ratio over 60 s bins, and log band power in four bands (0.5–4, 4–8, 15–30, 30–50 Hz) from a Welch spectrum of the spike train binned at 100 Hz over the first 900 s. Features are standardized on the training fold and classified by multinomial logistic regression (max_iter = 3000).
- **ISI-MLP**: the matched deep single-unit baseline from the same study^6^, using the 100-bin ISI histogram over 0–3 s, normalized to sum to one, as the only input. The classifier is a multilayer perceptron with hidden sizes (128, 64), ReLU activations and dropout 0.3 on both hidden layers, trained with Adam (learning rate 10*^−^*^3^, weight decay 10*^−^*^4^, batch size 4096, 60 epochs) on training-fold-standardized features.
- **Protocol shared by the two single-unit baselines**: both follow the evaluation protocol of the study they reproduce. The training fold is SMOTE-oversampled to class balance (with *k*_neighbors_ clamped to the rarest class) rather than class-weighted, and five seeds *×* five folds are run in two regimes. In the transductive regime, folds are stratified by region class, and in the inductive regime whole animals are held out. The inductive split is the one reported throughout.
- **NuCLR** ^14^: permutation-equivariant spatiotemporal transformer that embeds each unit from the surrounding population’s activity under a contrastive objective, re-run from its released code in a pinned environment. A source audit confirms the released IBL region model consumes only a binned spike-rate tensor. Its forward pass reads an array built solely from spike timestamps, and the model and dataset code contain no electrode position, depth or channel input. Following NuCLR’s recommendation to set the epoch count so that total training reaches *∼*50,000 steps, we train to a *∼*50,000-step budget at a starting batch size of 16, automatically halved on CUDA out-of-memory for the largest sessions (NuCLR’s per-batch population attention is *O*(units^2^) and exceeds the 40 GB A100 budget). On the IBL matched good-units corpus (62,856 QC-passing units, 693 insertions, 139 animals) this budget is reached at 10 epochs (*≈*39,200 steps) under five leave-animal-out splits, with the published 10 s contrastive view duration (a 10-latent temporal context at the model’s 1 s latent step). On the Allen CCF region tasks (super-region and the hippocampal formation, thalamus and visual areas sub-region partitions; Supplementary Fig. 23) the same *∼*50,000-step budget is met with per-task derived epoch counts, since the number of sessions carrying units for each task sets the steps per epoch. Our NuCLR re-run reaches macro-F1 0.453 against the 0.530 reported for NuCLR’s inductive zero-shot setting. The two protocols are both inductive but differ in how the held-out set is grouped. NuCLR’s zero-shot split holds out unseen sessions and probe insertions, whereas ours holds out whole animals, so with roughly five insertions per animal in this corpus an animal may contribute to both training and test under the published protocol but not under ours. That, together with the matched good-units corpus, is the most likely source of the gap.

## Data availability

All datasets used are public, including the Hull and Lisberger datasets via the C4 database^8^, the Buzsáki/CellExplorer dataset ^39^, the Allen Visual Coding corpus^4,16^, the IBL Brain Wide Map^17^, the forebrain-organoid HD-MEA recordings (DANDI:001374) ^25^, the Triple-N macaque ventral-stream recordings (Science Data Bank, https://doi.org/10.57760/sciencedb.33556)^23^, and the SUMMER human medial-temporal-lobe dataset (DANDI:001616, version 0.260702.0824, CC-BY-4.0) ^28^. Our NWB-converted datasets will be released with a citable archival DOI on publication.

## Code availability

All code is in a single repository (https://github.com/braingeneers/mycel-ephys), with a pinned environment specification, released under the Creative Commons Attribution-NonCommercial-NoDerivatives 4.0 International license (CC BY-NC-ND 4.0). It will be made public and archived on Zenodo with a citable DOI on publication.

## Funding Statement

This work was supported by Schmidt Futures (SF857) to M.T. and D.H.; the National Human Genome Research Institute (RM1HG011543) to M.T. and D.H.; the National Science Foundation (2134955) to M.T. and D.H., (2034037) to M.T., (2515389) to M.T., D.H., and M.A.M.-R.; the National Institute of Mental Health (U24MH132628) to D.H. and M.A.M.-R.; the National Institute of Neurological Disorders and Stroke (U24NS146314) to M.A.M.-R. and D.H.; the California Institute for Regenerative Medicine (DISC4-16285) to M.T. and M.A.M.-R., (DISC4-16337) to M.A.M.-R., and (DISC4-19334) to M.A.M.-R.; the University of California Office of the President (M25PR9045) to M.T. and M.A.M.-R.; and the Brain and Behavior Research Foundation (33184) to M.A.M.-R. J.G.-F. was supported by the QB3 Graduate Innovators Fellowship. The funders had no role in study design, data analysis, decision to publish, or preparation of the manuscript. The content is solely the responsibility of the authors and does not necessarily represent the official views of the National Institutes of Health, the National Science Foundation, the University of California, the California Institute for Regenerative Medicine, or any other federal or state agency.

## Acknowledgements

We thank the National Research Platform, supported by the NSF (CNS-1730158, ACI-1540112, ACI-1541349, OAC-1826967), the UC Office of the President, and UC San Diegos Qualcomm Institute.

During the preparation of this work, the authors used the Claude, Gemini, and ChatGPT large language models in order to generate portions of the source code and analysis figures. After using these tools, the authors reviewed and edited the content as needed and take full responsibility for the content of the publication.

## Author contributions

J.G.-F. designed the study and performed the experiments. A.M.-O. assisted with figure preparation. M.A.M.-R. supervised the study. A.S., J.R.M., M.T., and D.H. contributed to study design and interpretation. M.T., D.H. and M.A.M.-R. acquired funding. J.G.-F. and M.A.M.-R. wrote the manuscript, with contributions from all authors. All authors read and approved the manuscript.

## Competing interests

J.G.-F. and M.A.M.-R. are listed as inventors on provisional patent applications associated with the work presented in this manuscript. M.A.M.-R. is an advisor for Atoll Financial Group and Optimal, neither of which had any role in this study. The remaining authors declare no competing interests.

## Supplementary Information

**Supplementary Table 1:** MYCEL robustness controls on the matched IBL good-units set (leave-animal-out split, shape-and-timing node features and depth edges unless noted; macro-F1, mean *±* s.d. over *n* cross-validation splits, cv_seed 0–4, each an independently drawn animal partition).

| Control | Variant | $n$ | Macro-F1 |
| --- | --- | --- | --- |
| Message-passing backbone | GATv2 (ours, 2-layer) | 5 | $0.496 \pm 0.027$ |
| | GCN<sup>2</sup> | 3 | $0.492 \pm 0.018$ |
| | GraphSAGE<sup>3</sup> | 3 | $0.475 \pm 0.009$ |
| | GATv2 deep (3-layer) | 5 | $0.494 \pm 0.018$ |
| Network depth | 1 layer | 5 | $0.467 \pm 0.017$ |
| | 2 layers (baseline) | 5 | $0.496 \pm 0.027$ |
| | 3 layers | 5 | $0.494 \pm 0.018$ |
| Attention heads | 1 | 5 | $0.480 \pm 0.031$ |
| | 2 | 5 | $0.470 \pm 0.044$ |
| | 4 (baseline) | 5 | $0.496 \pm 0.027$ |
| | 8 | 5 | $0.492 \pm 0.042$ |
| Edge relation (in isolation) | co-firing only | 5 | $0.404 \pm 0.011$ |
| | cross-correlogram only | 5 | $0.410 \pm 0.012$ |
| Edge relation (added to depth) | depth + co-firing | 5 | $0.511 \pm 0.036$ |
| | depth + cross-correlogram | 5 | $0.476 \pm 0.020$ |

**Supplementary Table 2:** One-time cost of the encoding search. Wall time of the production NSGA-II run, summed over every recorded candidate evaluation (pass-1) and every full-budget retrain of the Pareto survivors (pass-2). “Hours” is aggregate evaluation time, the like-for-like quantity to compare against the single-run per-fold costs of Supplementary Fig. 9; “Elapsed” is the pass-1 end-to-end wall clock, which is shorter because up to 32 candidates are evaluated concurrently. The search is run once, on datasets of 103–668 units. The node representation it motivated is fixed once and used unchanged in every region, cross-species and cross-preparation experiment reported here, so the search is not repeated per task or per fold.

| Dataset | Pass-1 (low budget) |  | Pass-2 (full budget) |  | Total |  |
| --- | --- | --- | --- | --- | --- | --- |
|  | Evals | Hours | Retrains | Hours | Hours | Elapsed |
| Hull | 512 | 20.4 | 115 | 0.7 | 21.1 | 1.5 |
| Lisberger | 512 | 30.4 | 160 | 13.2 | 43.6 | 3.8 |
| Buzsaki | 512 | 28.3 | 135 | 37.5 | 65.8 | 2.5 |
| All three | 1536 | 79.1 | 410 | 51.4 | **130.5** | 7.8 |

**Supplementary Figure 1:**
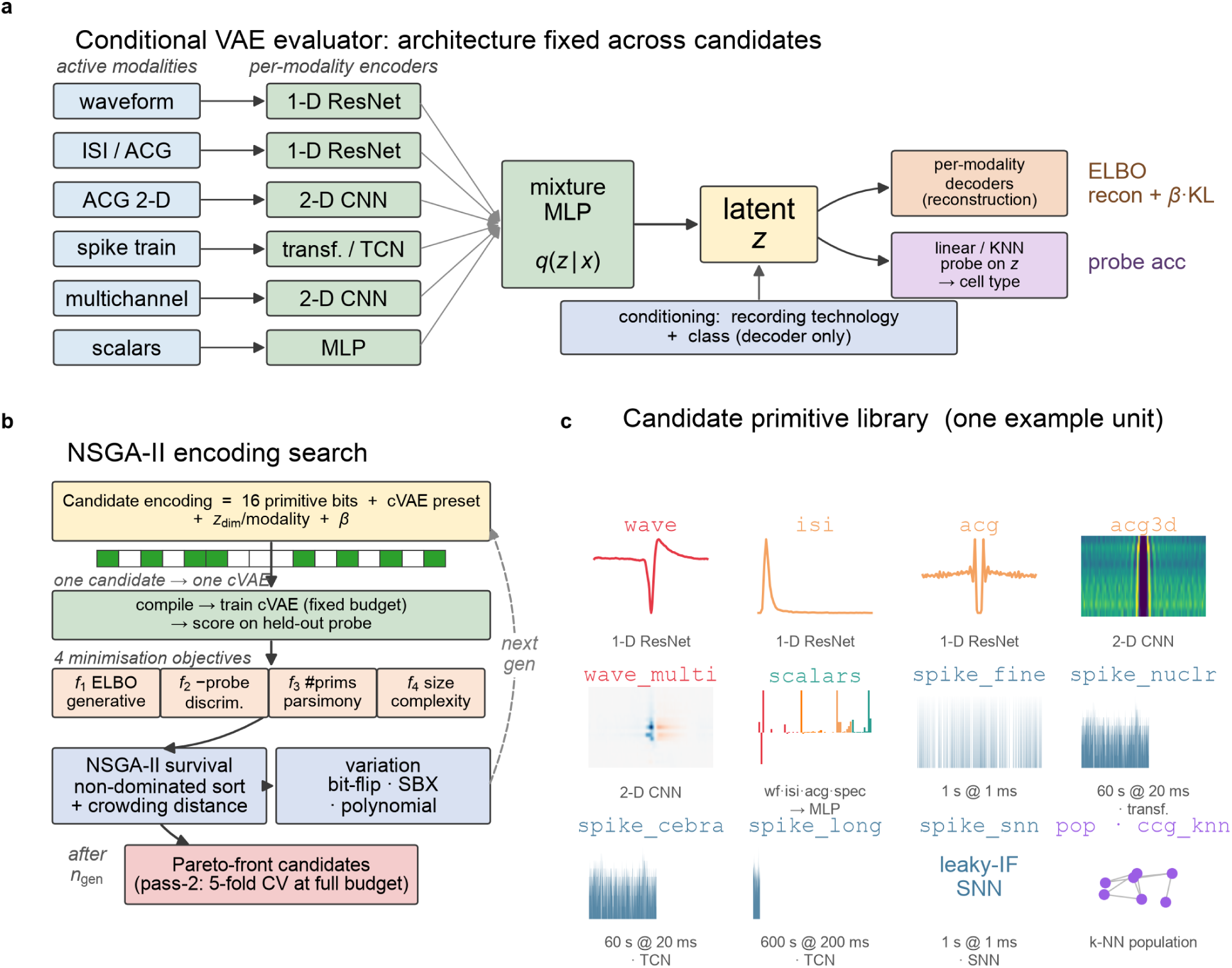
An evolutionary search over electrophysiological feature primitives. **a**, The cVAE evaluator, in which per-modality encoders feed a fusion multilayer perceptron (MLP) into a latent *z* conditioned on recording technology, with per-modality reconstruction decoders trained against an ELBO and a probe on *z* scoring cell-type accuracy; backbone family and training procedure are fixed across candidate encodings. **b**, NSGA-II search, in which each candidate encoding (16 primitive bits plus cVAE settings) compiles into a cVAE, is scored on four minimization objectives, and the surviving Pareto front seeds the next generation; survivors are retrained at full budget under five-fold cross-validation. **c**, The 16-primitive library shown on one example unit.

**Supplementary Figure 2:**
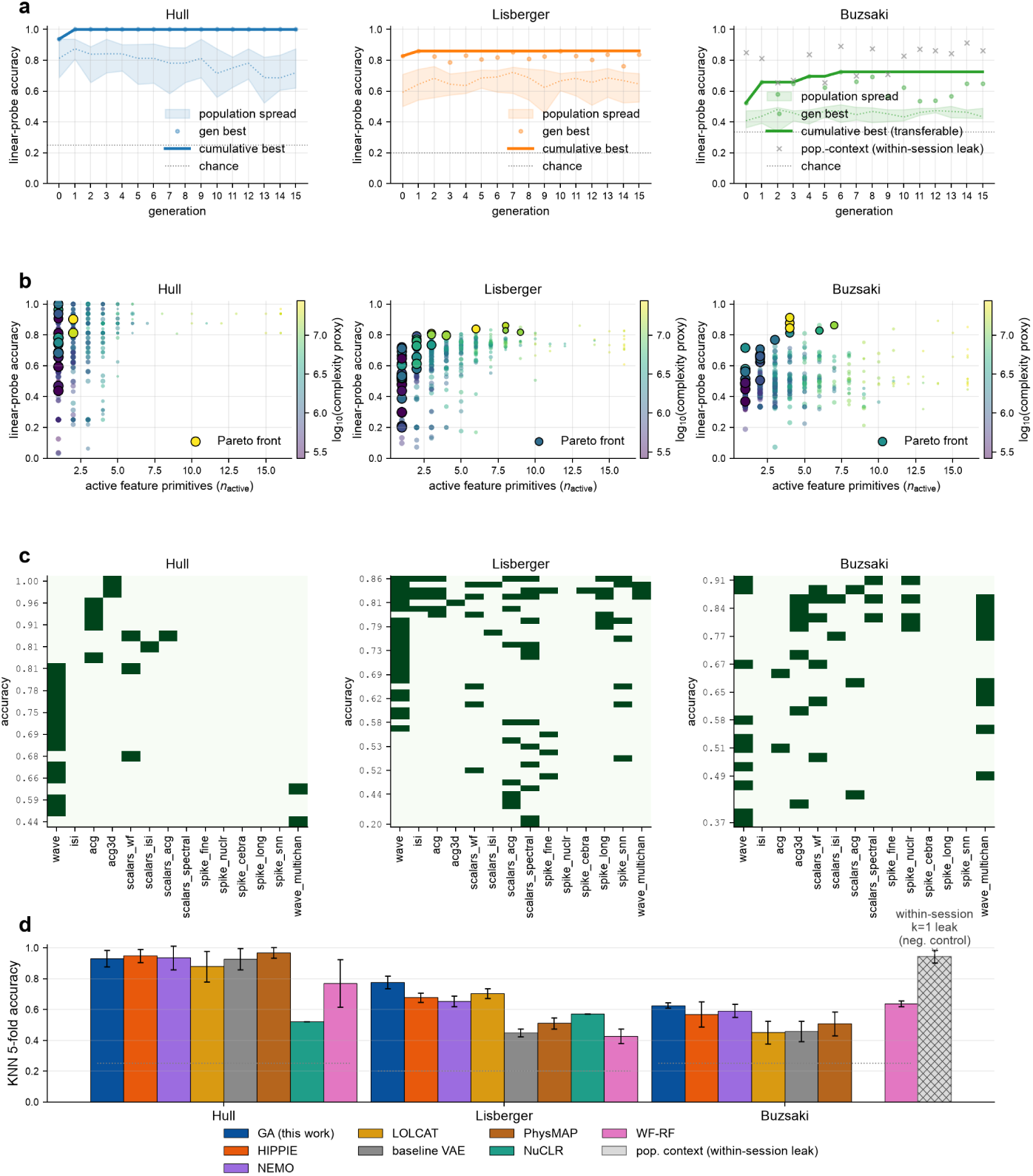
Evolutionary encoding search on three opto-tagged cell-type benchmarks. Columns in **a**–**c** are the Hull, Lisberger and Buzsáki datasets. **a**, Search progress over 16 NSGA-II generations, showing cumulative best-so-far balanced accuracy (bold) and per-generation best (markers). On Buzsáki the curve tracks transferable candidate encodings only, the population-context ones (gray *×*) being the within-session leak isolated in **d**. **b**, Pareto fronts in accuracy *×* active-primitive count, colored by log_10_ of the complexity proxy *f*_4_ (analytic, not measured FLOPs). **c**, Primitive composition of each front. **d**, Cross-validated balanced accuracy for the best candidate encoding (blue) versus baselines, each scored with its own published head (Methods); chance dotted (Hull 0.25; Lisberger 0.20; Buzsáki 0.25 on one common four-class interneuron taxonomy, 332 units). Hatched bar, negative control showing the population-context descriptor the search ranks first, a within-session *k*=1 lookup. NuCLR has no Buzsáki bar, its within-session contrastive objective being undefined on that dataset’s median 6-unit, class-pure sessions. Center, mean; error bars, s.d. over *n* = 5 folds. Lisberger, search-best versus HIPPIE, two-sided Welch’s *t*-test *P* = 0.003, Cohen’s *d* = 2.8; versus NEMO *P* = 0.001; versus LOLCAT *P* = 0.014, *d* = 2.0. For Hull, all tested baselines *P >* 0.08 (tie at ceiling); for Buzsáki, differences are not significant against the three strongest. Tests cover every baseline whose release reports per-fold scores. The search ran on fold 0; dropping it shifts the search-best means by at most 0.014.

**Supplementary Figure 3:**
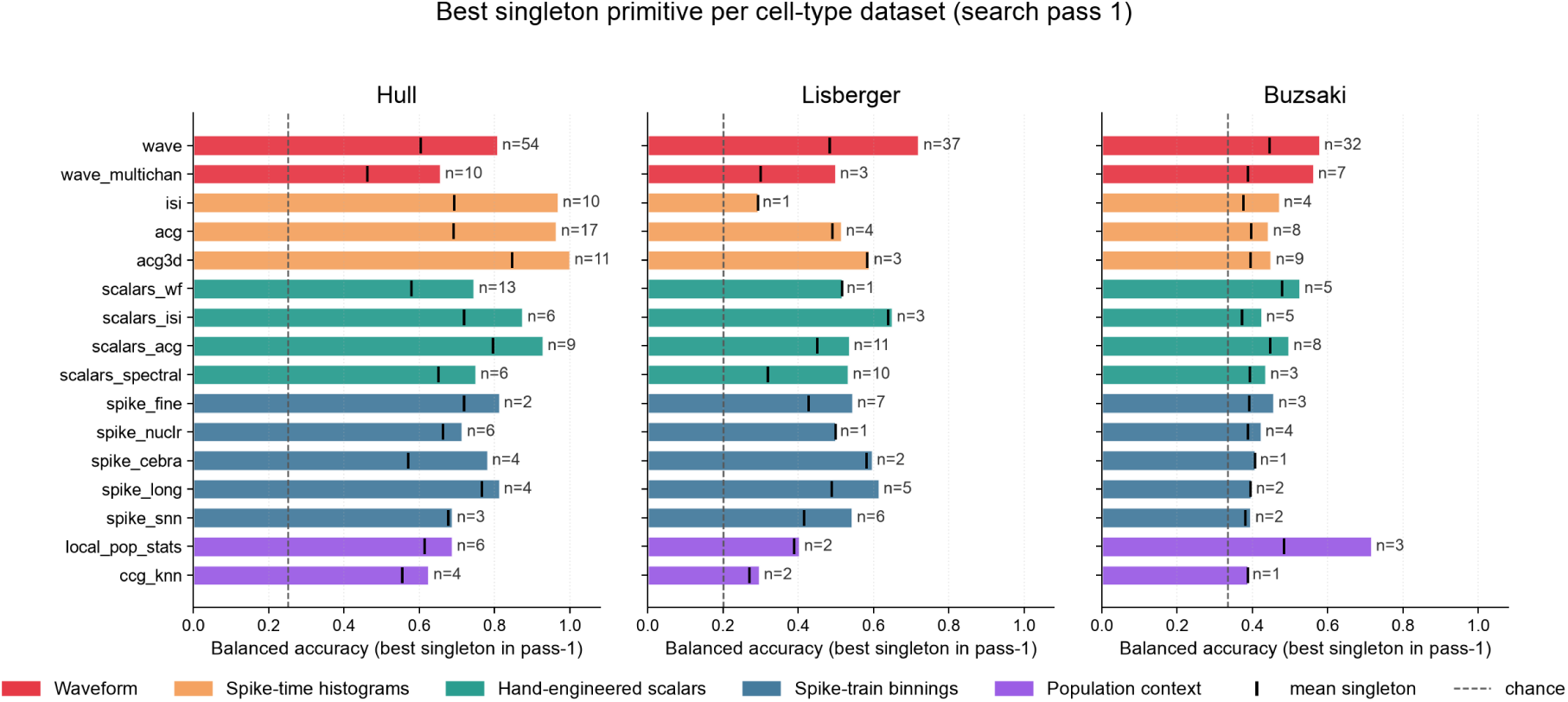
A different single primitive ranks highest on each cell-type dataset. Best singleton-primitive balanced accuracy per dataset (filled bar; *n* beside each bar is the number of singleton evaluations NSGA-II ran; black tick, mean; dashed line, chance). Chance differs by dataset because each singleton sweep uses that dataset’s own taxonomy, with Hull at 4 classes (0.25), Lisberger at 5 (0.20) and Buzsáki at 3 (0.33, the parvalbumin/somatostatin/vasoactive-intestinal-peptide default rather than the four-class taxonomy scored in Supplementary Fig. 2d). Hull is topped by the two-dimensional autocorrelogram, Lisberger by the peak waveform, Buzsáki by the population descriptor (the within-session shortcut quantified in the Methods, not a transferable cell-type feature). The optimal encoding therefore does not transfer, even at single-primitive granularity.

**Supplementary Figure 4:**
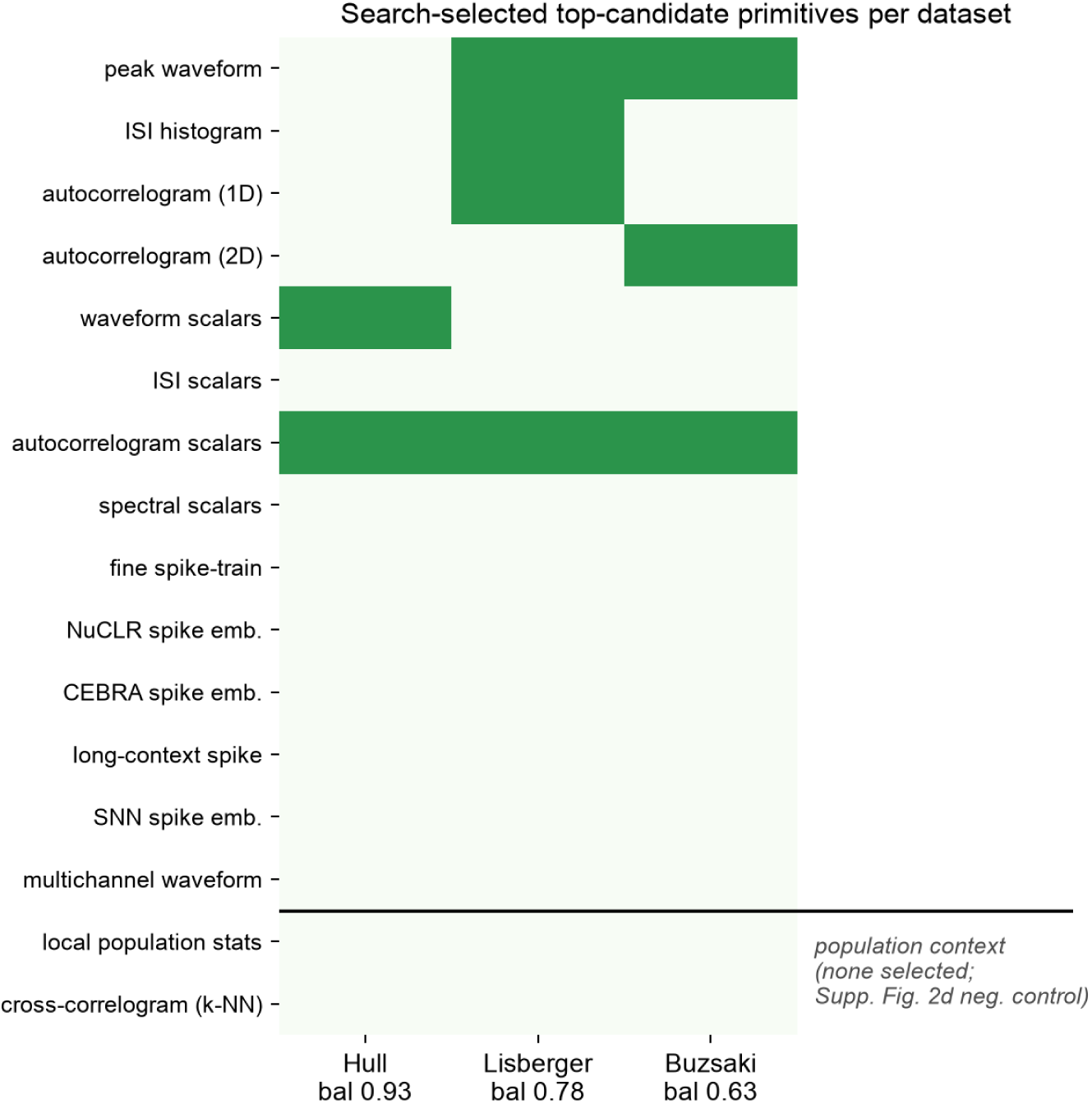
Primitive composition of the search-selected top candidate on each dataset. A filled cell marks a primitive active in that dataset’s top candidate (column header, 5-fold balanced accuracy). The two population-context primitives are drawn in a separate block, and neither is selected in any transferable top candidate (the Buzsáki population-context descriptor is the Supplementary Fig. 2d negative control). The autocorrelogram-scalar block is the only primitive selected on all three datasets.

**Supplementary Figure 5:**
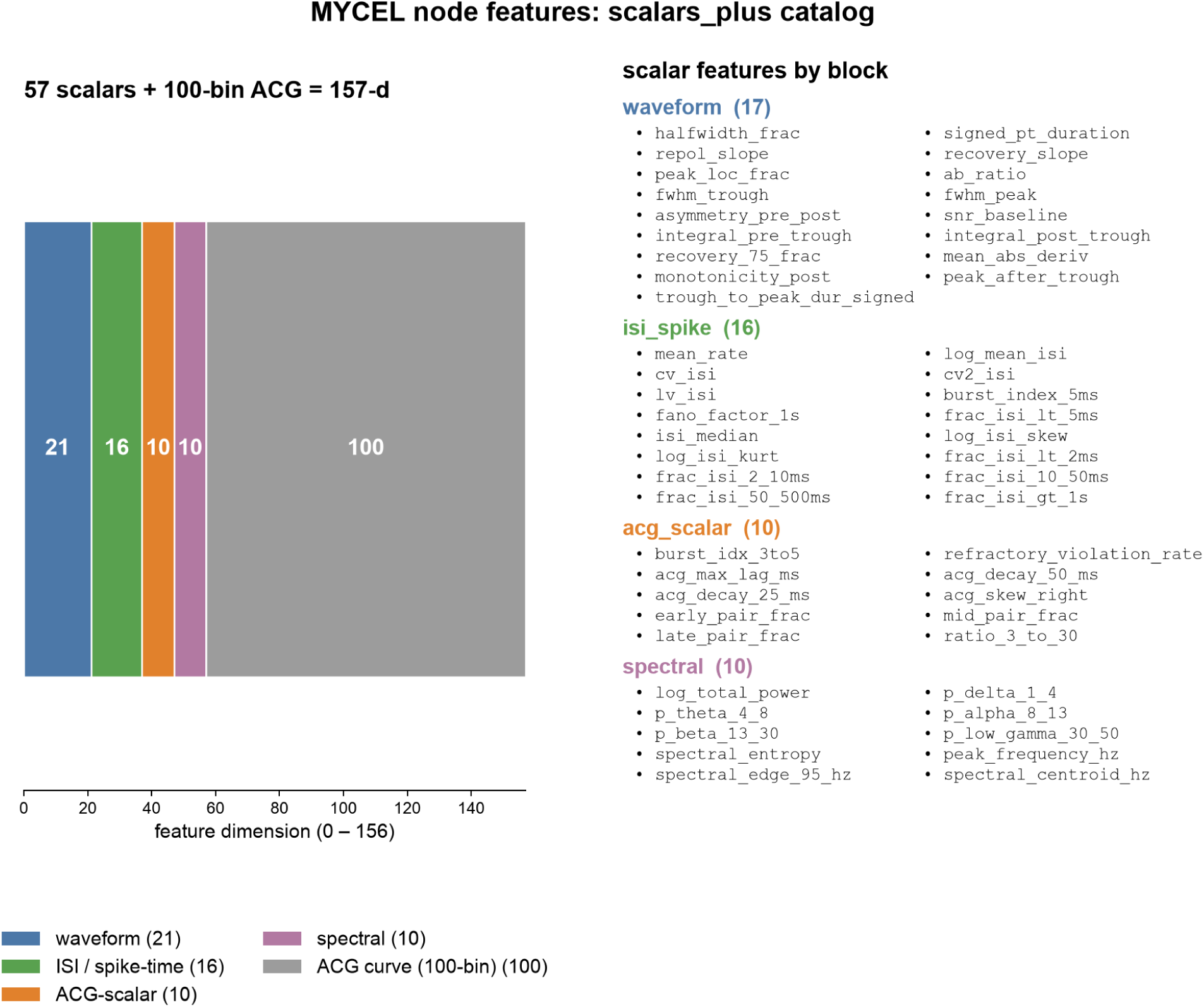
The shape-and-timing node features carried into the brain-region analysis. The 157-dimensional per-unit node representation used by MYCEL concatenates the four hand-engineered scalar blocks, waveform (21), ISI/spike-time (16), autocorrelogram (10) and spectral (10), 57 scalars total, with the 100-bin one-dimensional autocorrelogram histogram. Left, dimension budget per block; right, the individual scalar features by block (four constant-amplitude dimensions omitted). This is the feature set whose composition is dissected in the region results.

**Supplementary Figure 6:**
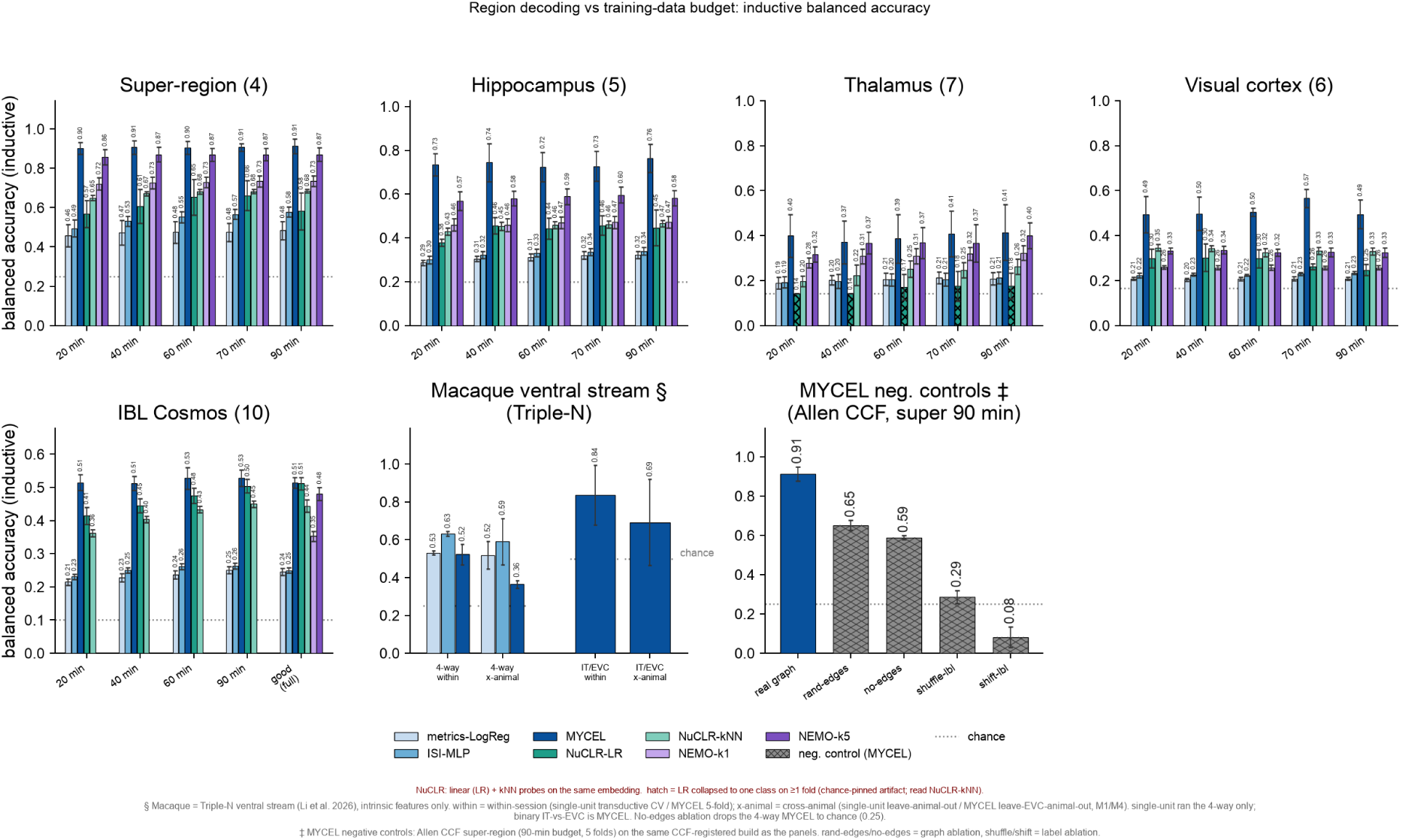
Full region-decoding benchmark across methods, tasks and recording-time budgets. Inductive (leave-animal-out) balanced accuracy for every method, the single-unit encoders (metrics-LogReg, ISI-MLP), MYCEL, NuCLR with its linear (LR) and *k*NN probes, and NEMO at *k*=1 and *k*=5 (at every Allen budget; on IBL Cosmos at full recording time only), as a function of the per-unit recording-time budget (20–90 min), on the five region tasks summarized in Fig. 2a, namely the Allen four-way super-region and its hippocampal formation (5-way), thalamus (7-way) and visual areas (6-way) sub-region partitions, and the IBL ten-way Cosmos partition (good units, with the full-recording point at right). The two rightmost panels add the macaque Triple-N ventral stream^1^ (4-way and binary IT-versus-EVC, within-session and cross-animal) and the MYCEL negative controls on the Allen super-region (real graph, random edges, no edges, label-shuffle, label-shift). Dotted line, chance; hatched NuCLR bars mark folds where its linear probe collapses to one class (read the NuCLR-*k*NN bar there).

**Supplementary Figure 7:**
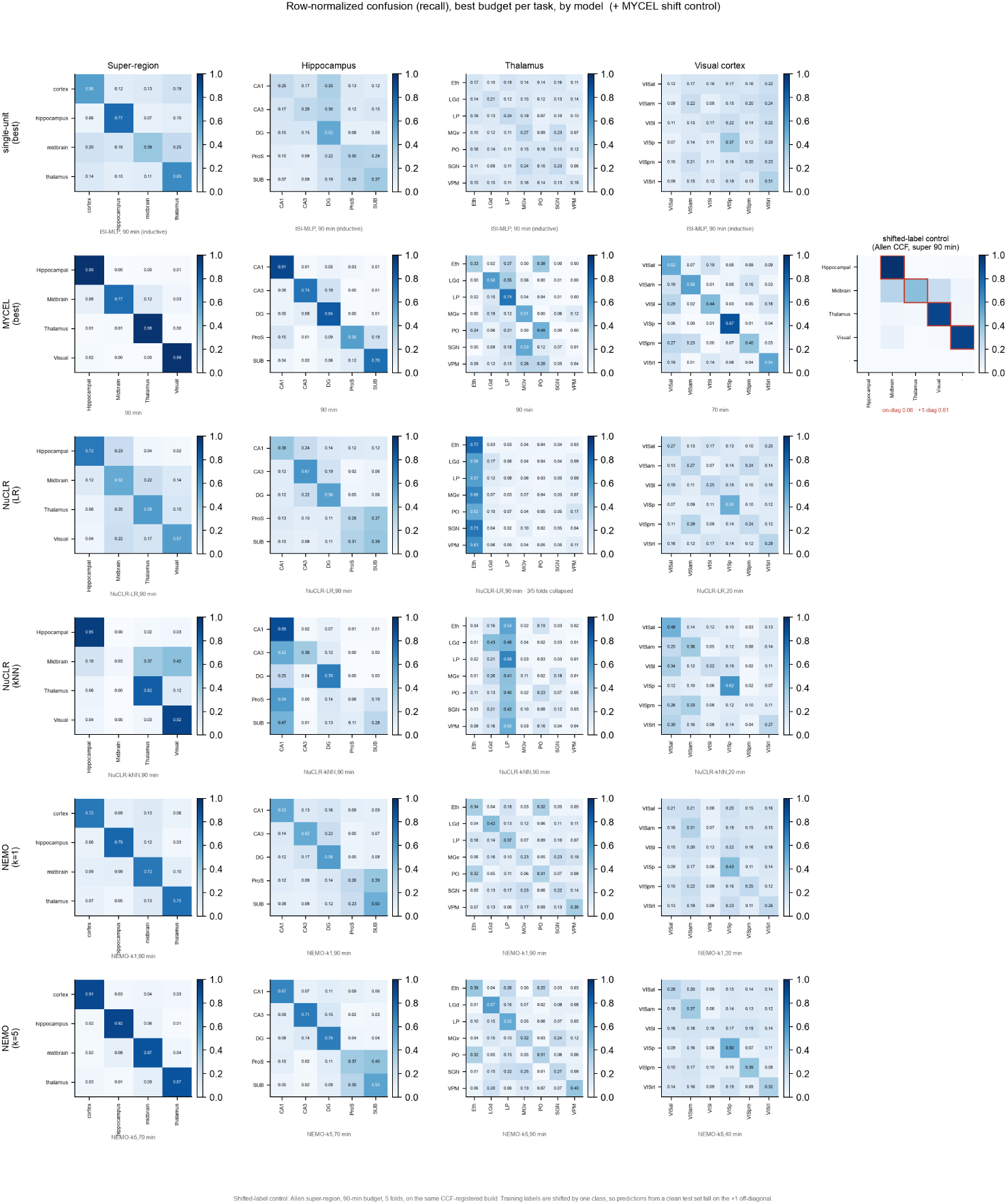
Which regions are confused for which, across Allen tasks. Row-normalized confusion (recall) on the CCF-registered Allen build, one panel per decoding task (super-region and the hippocampal formation, thalamus and visual areas sub-region partitions) and one row per method. Counts are summed over the inductive (leave-animal-out) folds and then row-normalized, the same split the balanced accuracies in Fig. 2 and Supplementary Fig. 6 are computed on. Each row is shown at the budget where that method scores highest on that task, labeled per panel. Right of the MYCEL row, the label-shift control on the super-region (90 min). The single-unit row shows whichever of metrics-LogReg and ISI-MLP scores higher on that task, named in the panel subtitle. Diagonal mass reflects the recovered anatomy, and off-diagonal entries identify the neighboring areas that are hardest to separate.

**Supplementary Figure 8:**
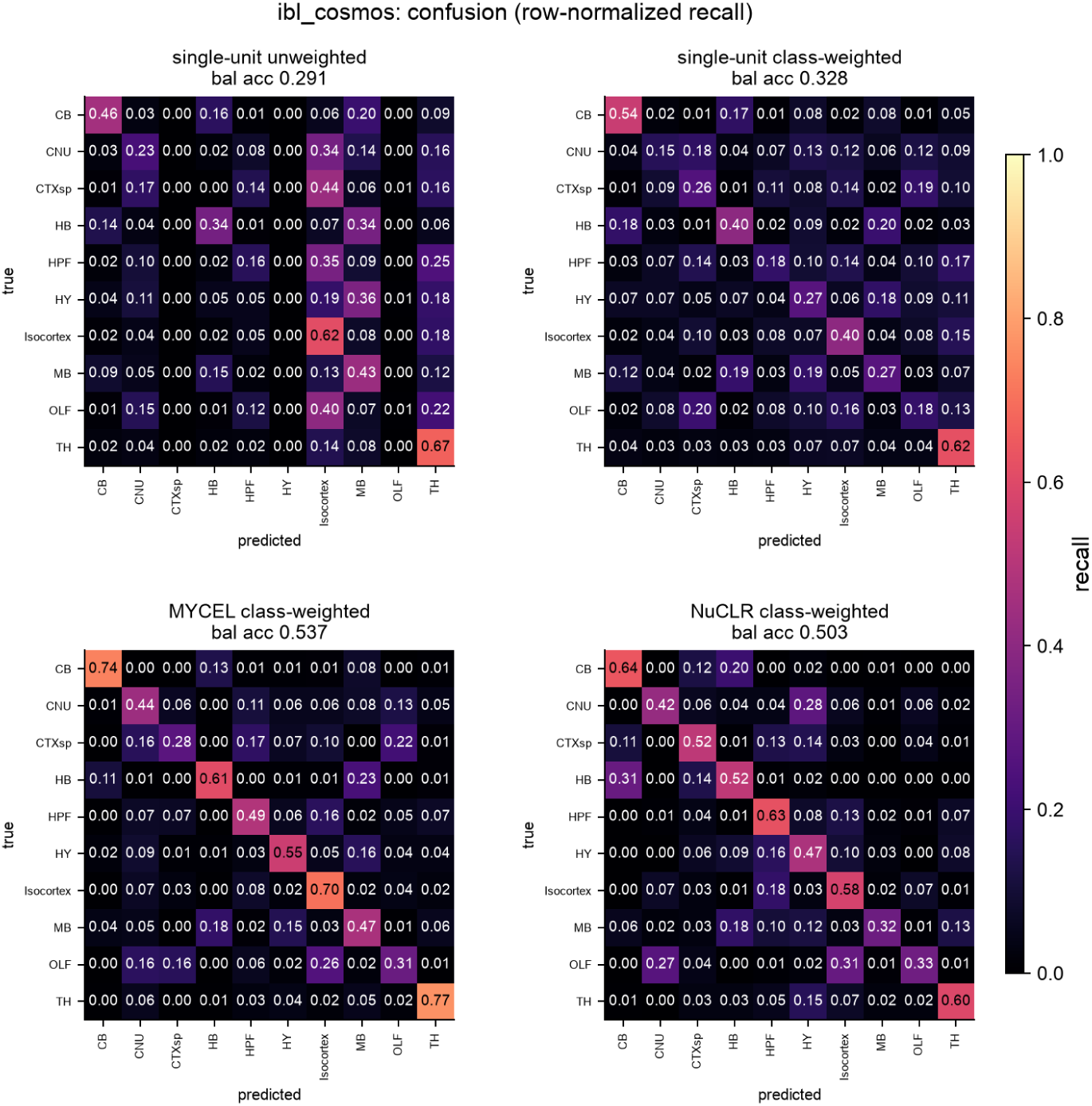
On IBL Cosmos the population graph resolves regions that single units smear together. Row-normalized confusion (recall) on the 10-region Cosmos taxonomy, matched good units. Single-unit scalars (top; unweighted then class-weighted) stay diffuse even when weighted (balanced accuracy 0.291 *→* 0.328), whereas the class-weighted MYCEL (bottom-left) shows a sharp diagonal (Cerebellum 0.74, Isocortex 0.70, Thalamus 0.77; balanced accuracy 0.537) and NuCLR (bottom-right) reaches 0.503, both well above the single-unit baseline.

**Supplementary Figure 9:**
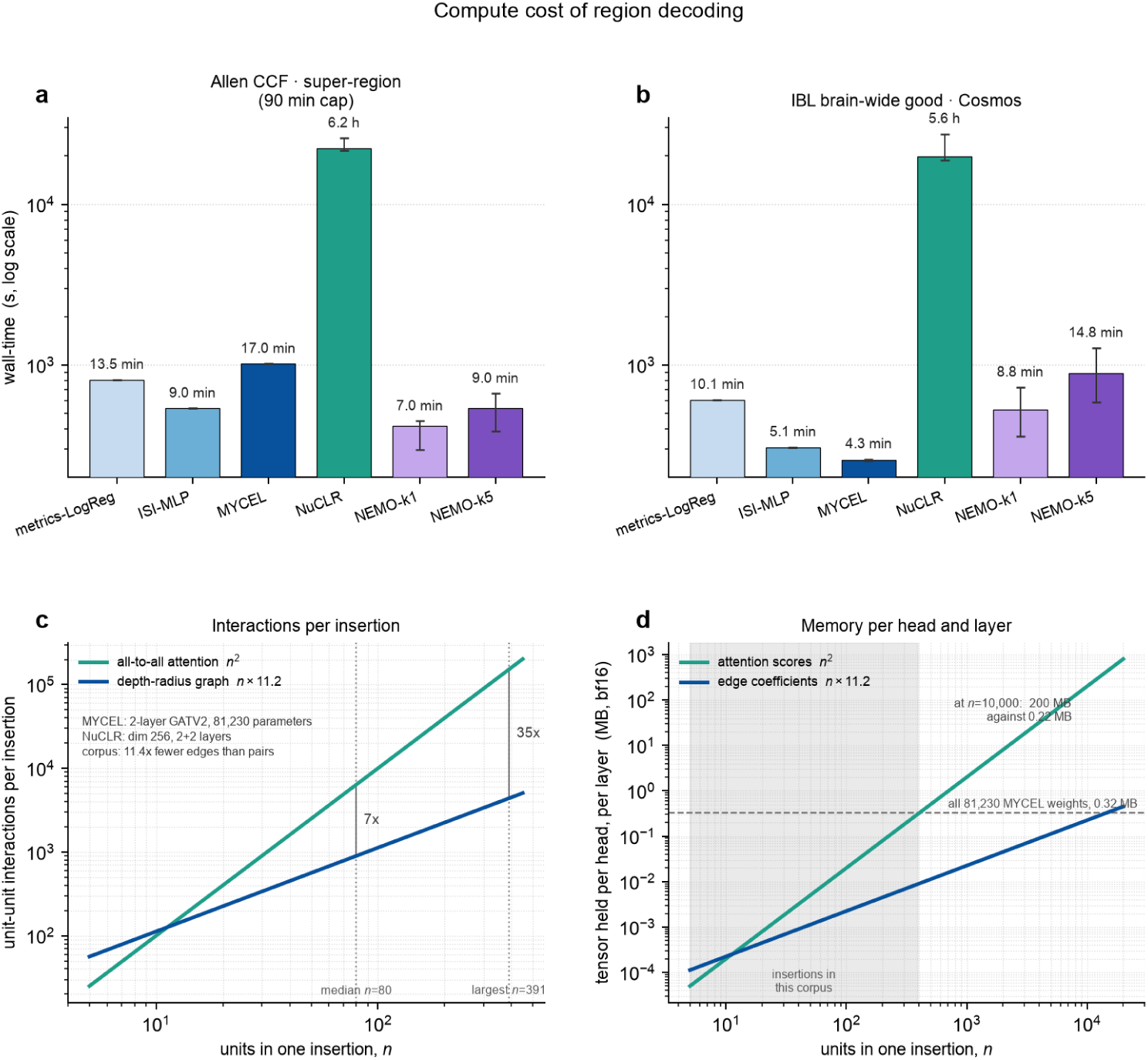
Compute cost of region decoding. **a**,**b**, Median wall time per fold (log scale; whiskers, range across folds) on the Allen super-region (90-min cap) and on IBL Cosmos good units, the two tasks on which all five methods were timed. MYCEL takes 17 min per fold on the Allen task and 4.3 min on IBL Cosmos, against 6.2 h and 5.6 h for NuCLR, whose cost is set by its *∼*40–50k-step training budget rather than by the size of the dataset. MYCEL ran on an 8 GB GPU and NuCLR on a *≥*40 GB GPU. The one-time encoding search (130.5 h; Supplementary Table 2) is not included. **c**, Unit-to-unit interactions per insertion of *n* units, *n*^2^ for all-to-all attention and 11.2*n* for the 200 *µ*m depth graph. Dotted lines mark the median (*n* = 80) and largest (*n* = 391) insertions, where attention computes 7 and 35 times as many. **d**, Memory for one head and one layer in bf16, comparing attention scores (*n*^2^) against edge coefficients (11.2*n*). Shading, the insertion sizes in this corpus. At *n* = 10,000 the attention scores take 200 MB and the edge coefficients 0.22 MB, less than the 0.32 MB of all 81,230 MYCEL weights (dashed line).

**Supplementary Figure 10:**
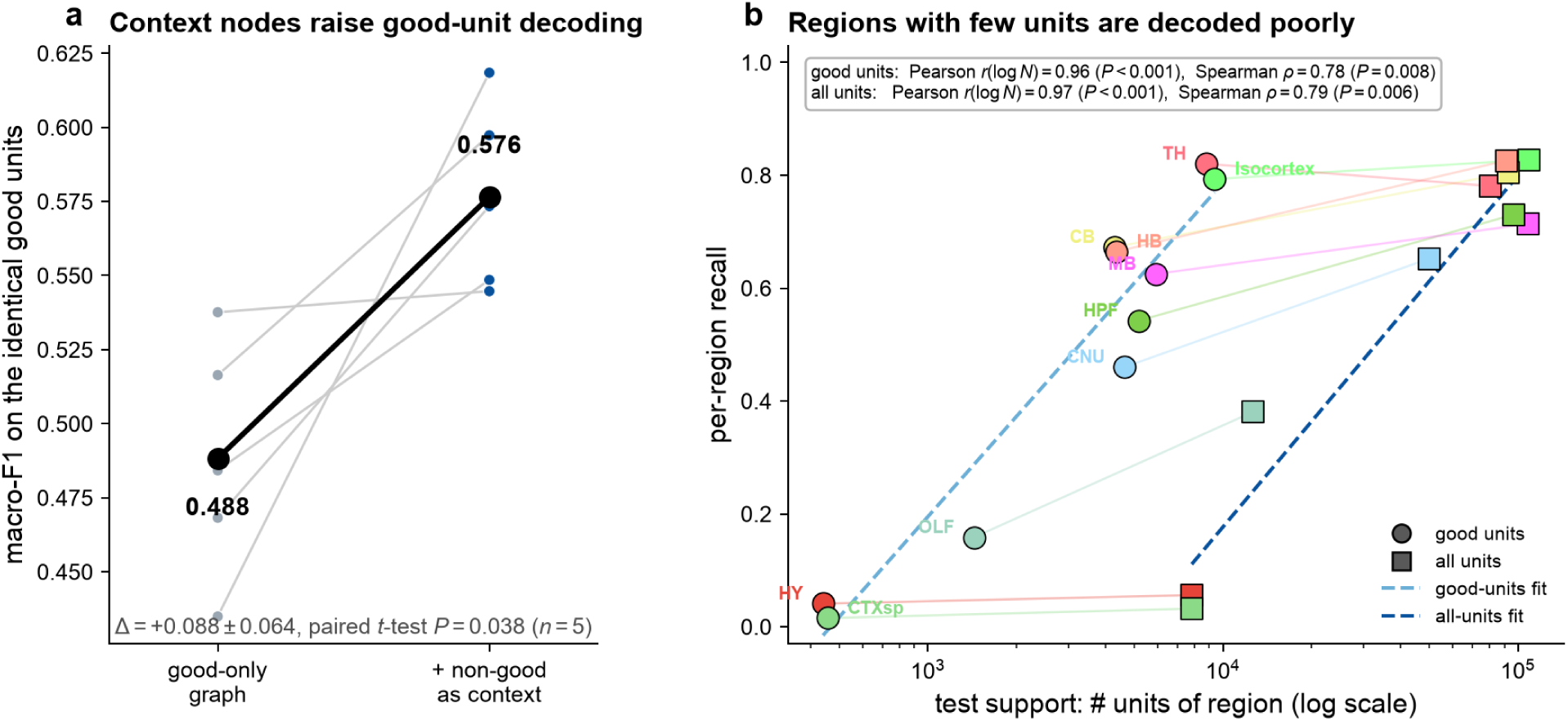
Units that fail quality control, kept as context-only nodes, improve good-unit decoding. MYCEL was trained on the full unit graph but scores only the QC-passing good units, so the non-good units act as context-only message-passing neighbors; metrics are on the *identical* scored good units as the good-only baseline (IBL Cosmos leave-animal-out split). **a**, Paired macro-F1 across the five inductive splits, good-only graph (0.488) versus all-units graph with good units scored (0.576; Δ = +0.088 *±* 0.064, paired *t*-test *P* = 0.038, higher in all five). Gray, per-split; black, mean. **b**, Per-region recall against the number of units the region supplies (test support; log scale, colored by the Allen ontology), for both MYCEL unit sets, good units (circles) and all units (squares), joined per region. Recall rises with unit supply under either set (good units: Pearson *r*(log *N*) = 0.96, Spearman *ρ* = 0.78, *P* = 0.008; all units: *r*(log *N*) = 0.97, *ρ* = 0.79, *P* = 0.006; dashed, log-support fits). Regions with many units (thalamus, isocortex) reach *∼*0.8, while regions with few units (hypothalamus, cortical subplate) fall below 0.15.

**Supplementary Figure 11:**
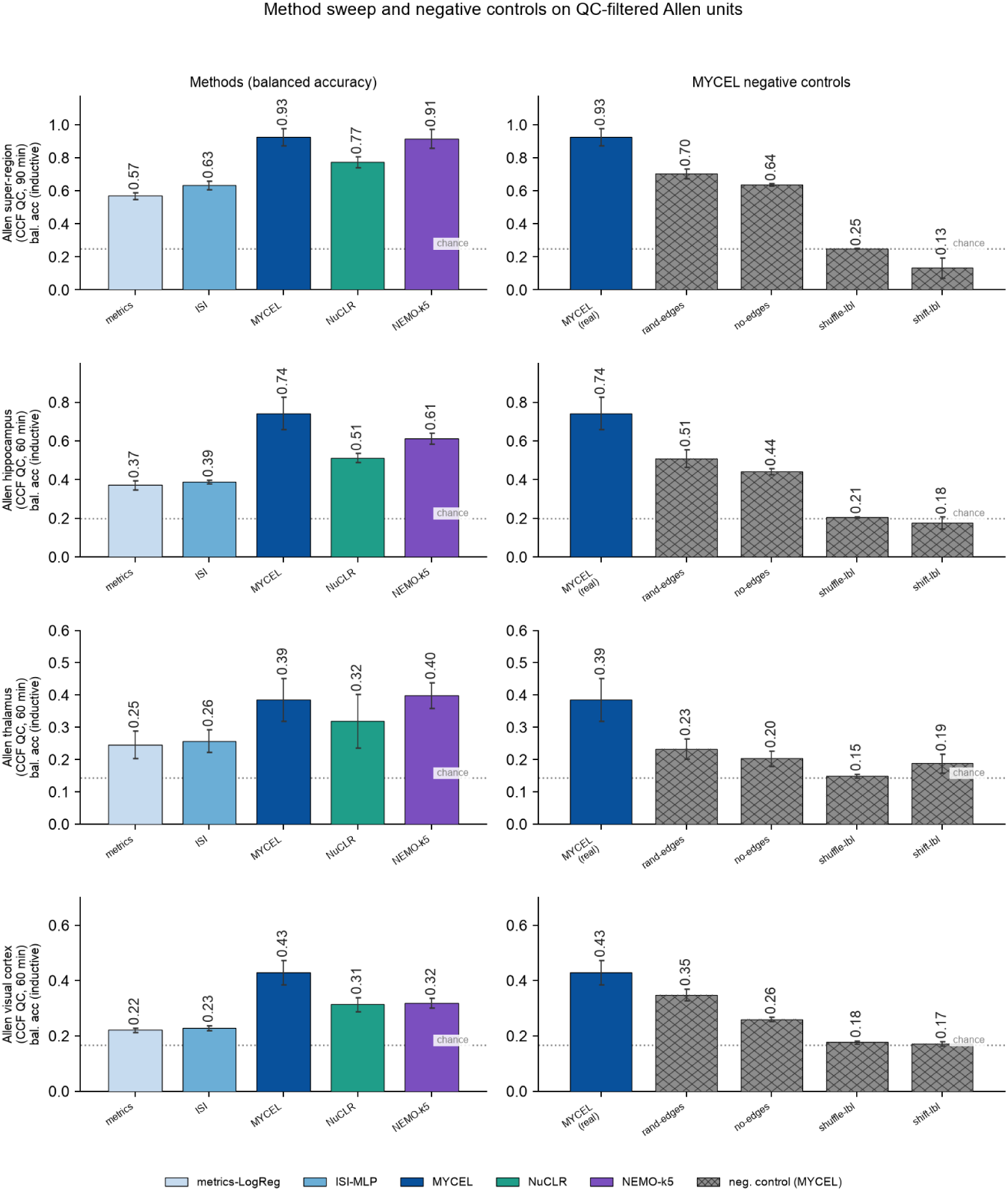
Method sweep and negative controls on QC-filtered Allen units. The sweep of Supplementary Fig. 23 repeated on the four Allen CCF tasks after the three AllenSDK default unit filters (ISI violations *<* 0.5, amplitude cutoff *<* 0.1, presence ratio *>* 0.9), with the same folds. MYCEL scores above NuCLR and both single-unit encoders on every task (super-region 0.93, hippocampal formation 0.74, thalamus 0.39, visual areas 0.43), and the label-shuffle and label-shift controls fall to near chance, so the decode does not depend on low-quality units. NEMO matches MYCEL on the super-region (0.91) and thalamus (0.40) but not on the hippocampal formation (0.61) or visual areas (0.32) partitions. Bars, per-fold mean; whiskers, s.d.

**Supplementary Figure 12:**
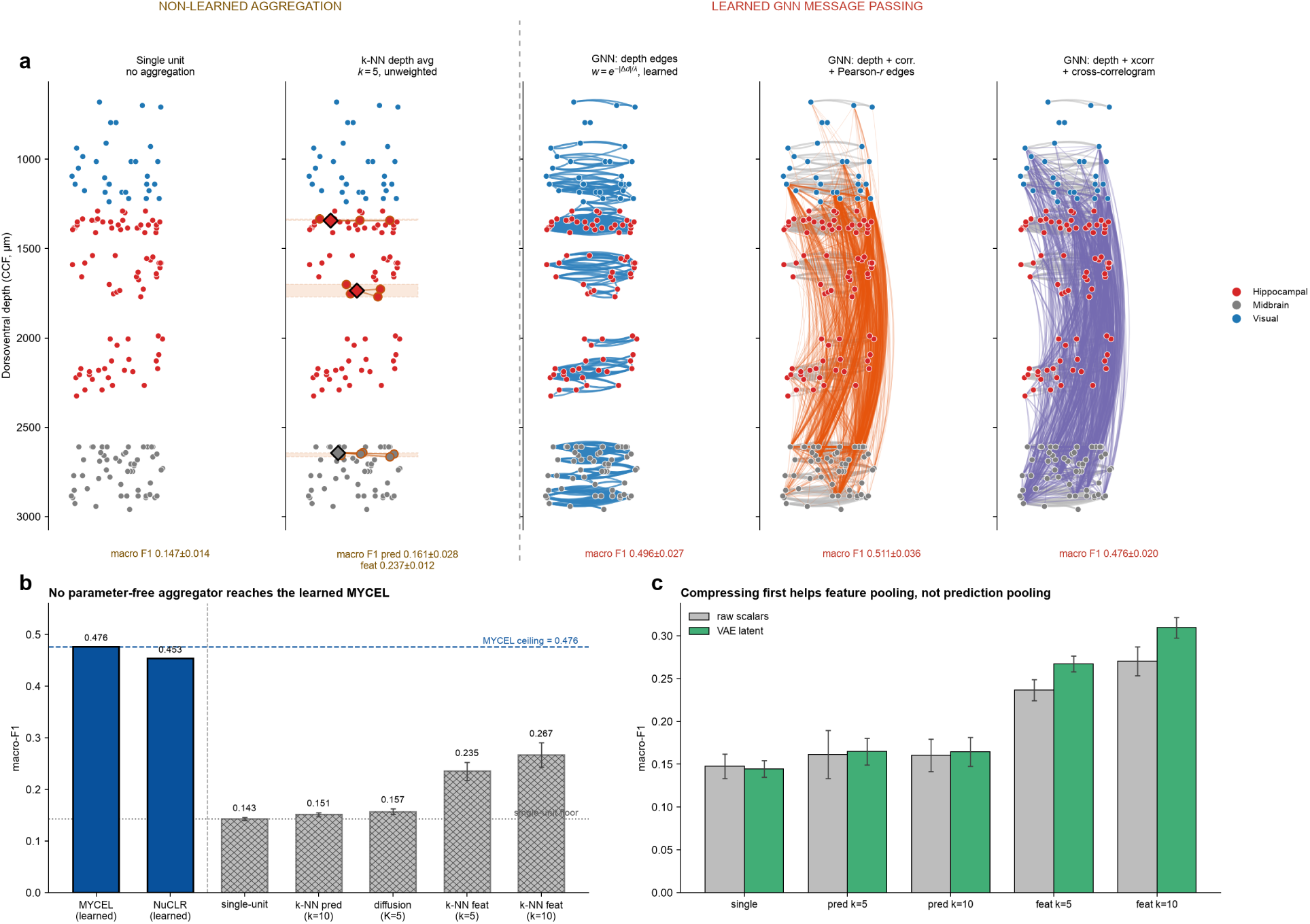
From no aggregation to the learned graph, and the parameter-free controls. IBL Cosmos, matched good units, five leave-animal-out splits (mean *±* s.d.), unless noted. **a**, Different aggregation methods represented on an example IBL Neuropixels insertion. Over the *identical* depth neighborhood, parameter-free aggregators (single-unit 0.147; prediction-average 0.161; feature mean-pool 0.237) sit far below the learned GATv2 (0.496 depth; 0.511 depth+co-firing); macro-F1 values are the method’s matched good-units five-split scores, not the single illustrative probe. The aggregators are at *k*=5; an exploratory *k* sweep peaks near *k*=20 and over-smooths beyond it, still far below the learned graph. **b**, Parameter-free aggregators quantified against both learned references, read from a separate sweep of the same configurations (MYCEL 0.476, NuCLR 0.453). Every parameter-free aggregator over the depth neighborhood sits well below the learned baselines. **c**, Replacing the raw shape-and-timing node features with a frozen unsupervised VAE latent (dimension 32) before the same cheap aggregators barely changes the single-unit and prediction-average scores, but gives a modest, reproducible gain to feature-pooling (*k*=5: 0.237 *→* 0.267; *k*=10: 0.271 *→* 0.310).

**Supplementary Figure 13:**
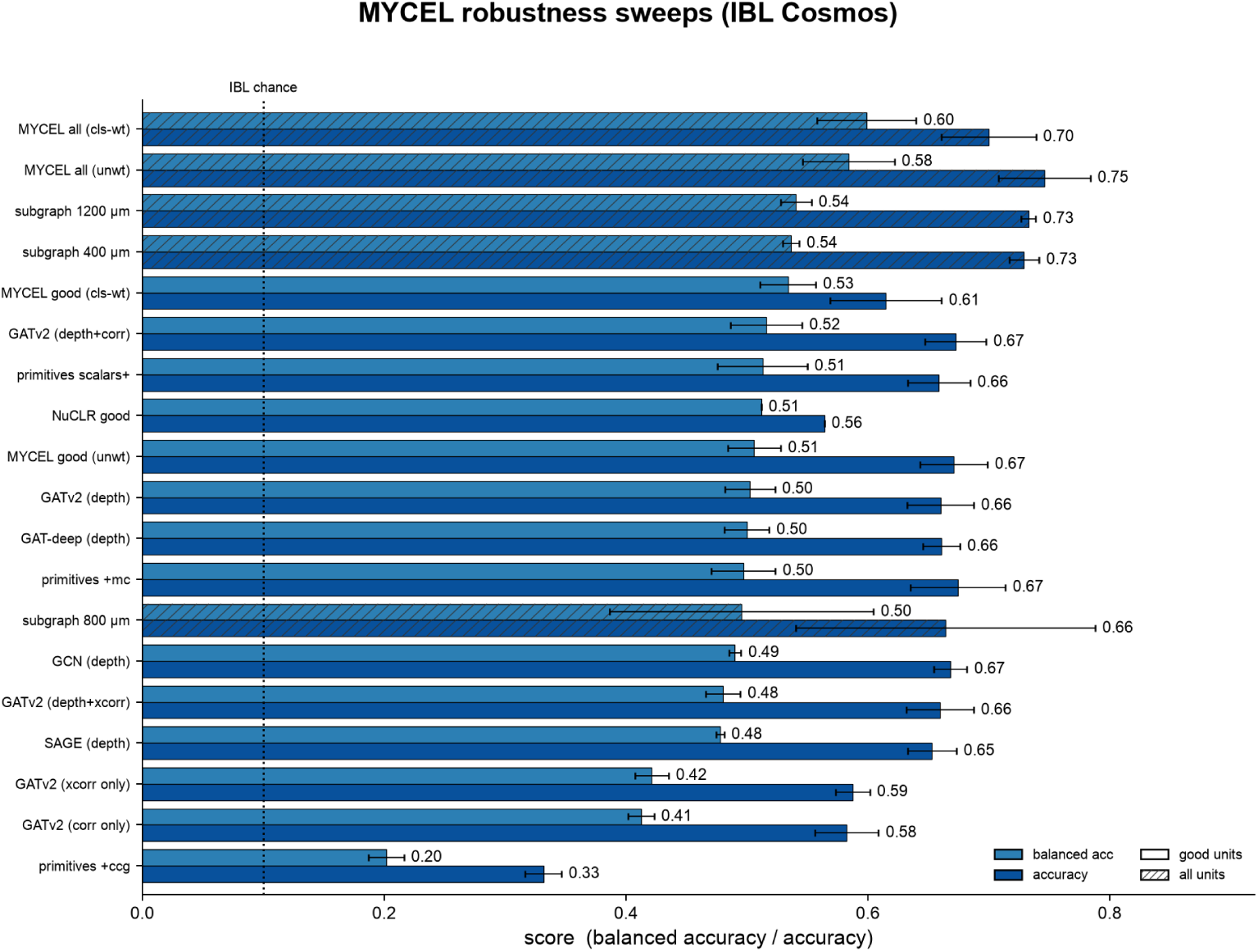
MYCEL robustness sweeps on IBL Cosmos. Balanced accuracy and accuracy across message-passing backbone (GCN / SAGE / GATdeep / GATv2), edge relation (depth *±* correlation / cross-correlogram, correlogram-only), node set, subgraph window and class-weighting, on good units (unhatched) or all units (hatched). Backbone, subgraph-window and class-weighting variants sit within a narrow band; correlogram-only edges and the cross-correlogram node set score lower. Center, mean; error bars, *±*1 s.d. over inductive splits or seeds (NuCLR, single run).

**Supplementary Figure 14:**
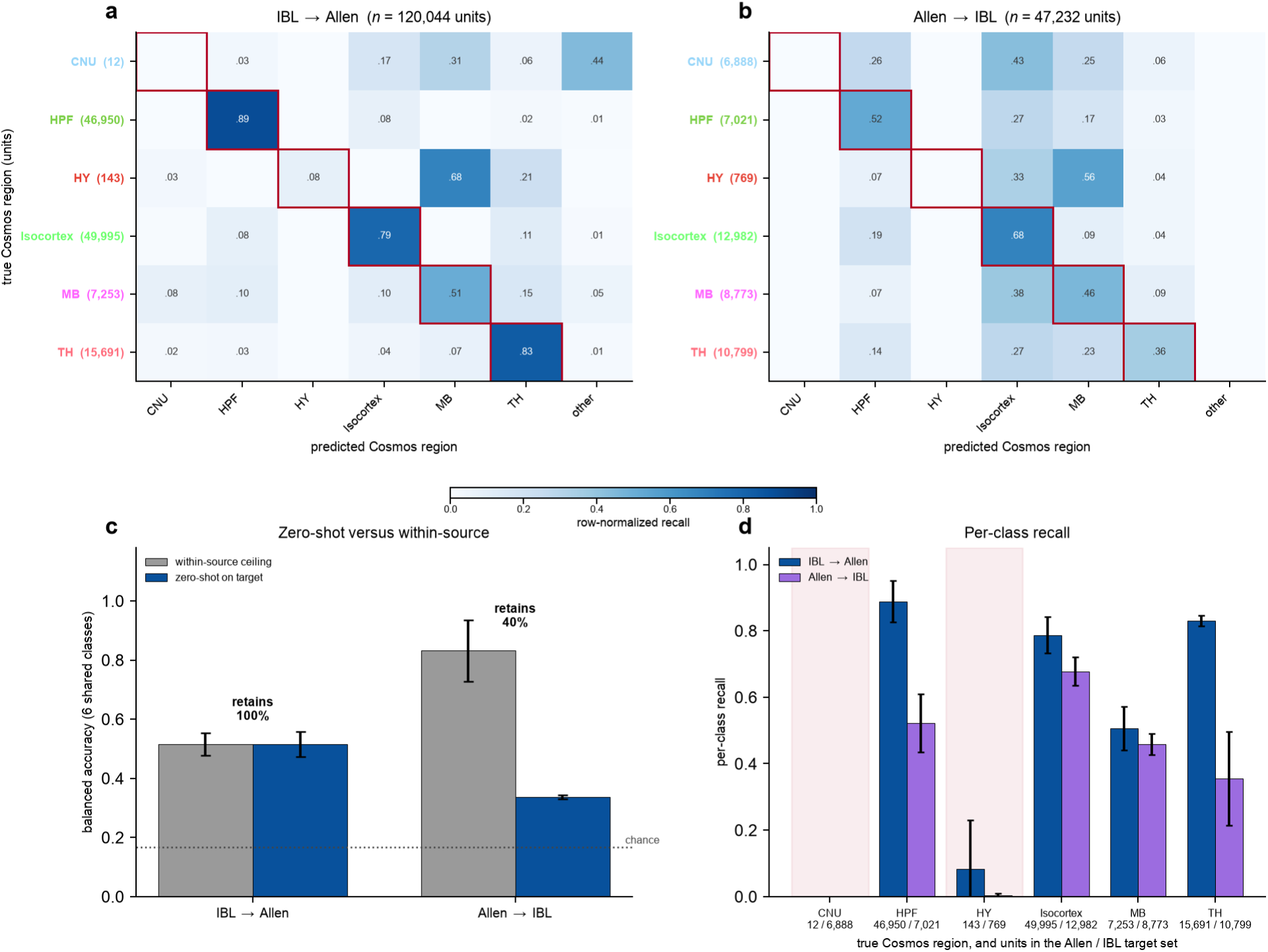
Cross-atlas zero-shot region transfer. MYCEL trained on one corpus and applied to the other without adaptation, scored on the six Cosmos classes the two corpora share (chance 0.167; *n* = 3 seeds per direction, mean *±* s.d.; Methods). **a**,**b**, Row-normalized recall for IBL*→*Allen (120,044 target units) and Allen*→*IBL (47,232). Red outline, correct class; “other”, predictions outside the shared classes. **c**, Zero-shot balanced accuracy against each model’s accuracy on held-out animals of its own corpus. **d**, Per-class recall in each direction, with the units per class in each target set. Shading, the two classes Allen Visual Coding barely samples.

**Supplementary Figure 15:**
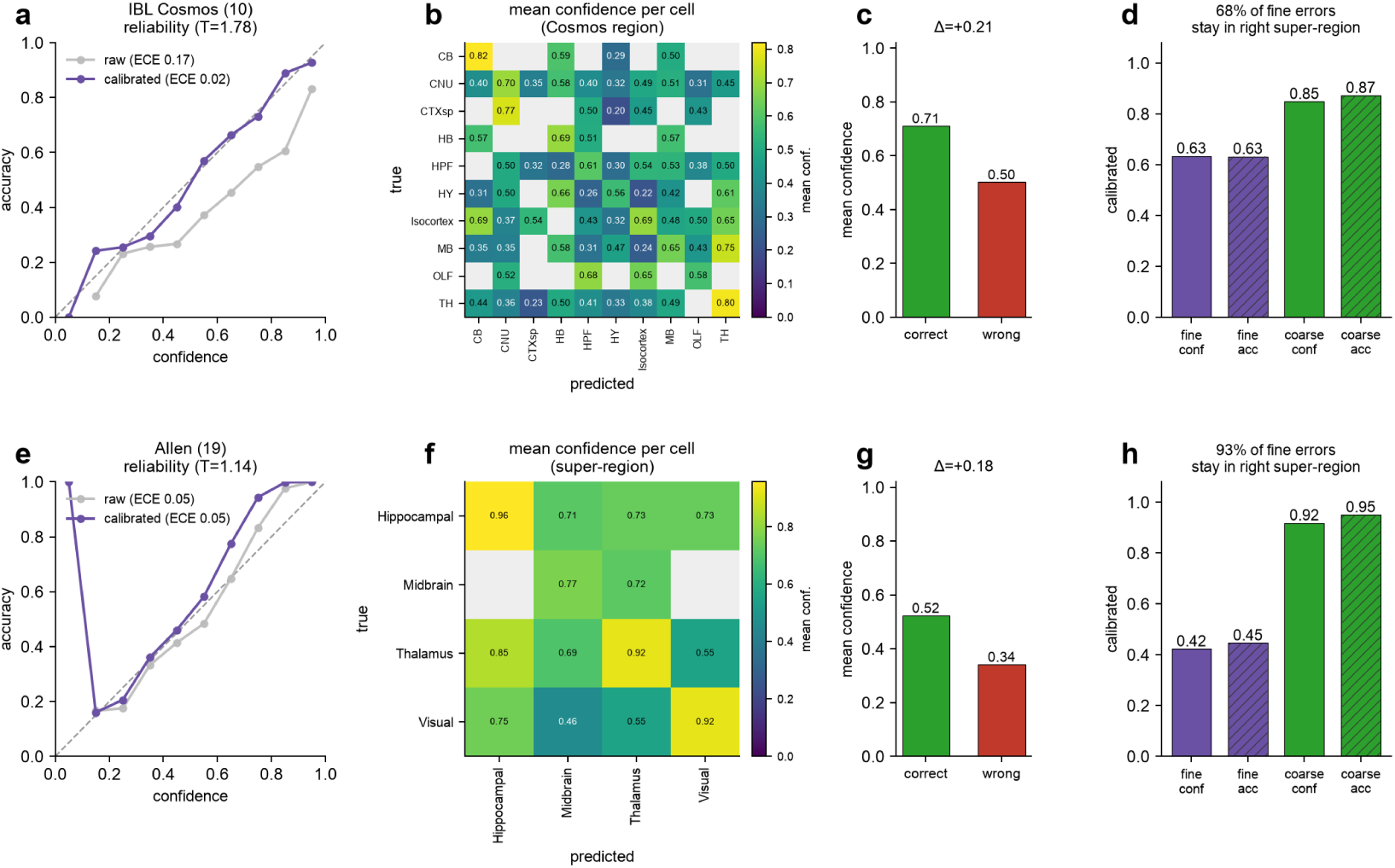
MYCEL’s region predictions are calibrated, error-aware and hierarchically usable. Every prediction carries a confidence score, the temperature-scaled max-softmax (Methods), for the two region tasks with an anatomical hierarchy; one row per task (top, IBL Cosmos, **a**–**d**; bottom, Allen, **e**–**h**). *T* is fit on an animal-disjoint validation split; every metric is on the untouched test split. **a**,**e**, Reliability diagrams (dashed, perfect calibration). IBL logits are over-confident (*T* = 1.78) and scaling cuts expected calibration error from 0.166 to 0.025; Allen is already calibrated (*T* = 1.14, ECE 0.049). **b**,**f**, Mean confidence per (true, predicted) cell (IBL at ten Cosmos regions; Allen at four super-regions; cells with *<* 3 predictions grayed). On average the diagonal is brighter than the off-diagonal, so the model is less confident when it errs. **c**,**g**, Mean confidence for correct versus wrong calls (IBL 0.71 vs 0.50, Allen 0.52 vs 0.34). **d**,**h**, Summing the fine softmax over each super-region’s children gives a calibrated coarse prediction. Coarse accuracy far exceeds fine (0.87 vs 0.63 IBL; 0.95 vs 0.45 Allen) at matched calibration, and 68% (IBL) and 93% (Allen) of fine errors still land in the correct super-region. Solid bars, confidence; hatched, accuracy. The binary organoid and macaque tasks are single-level and omitted.

**Supplementary Figure 16:**
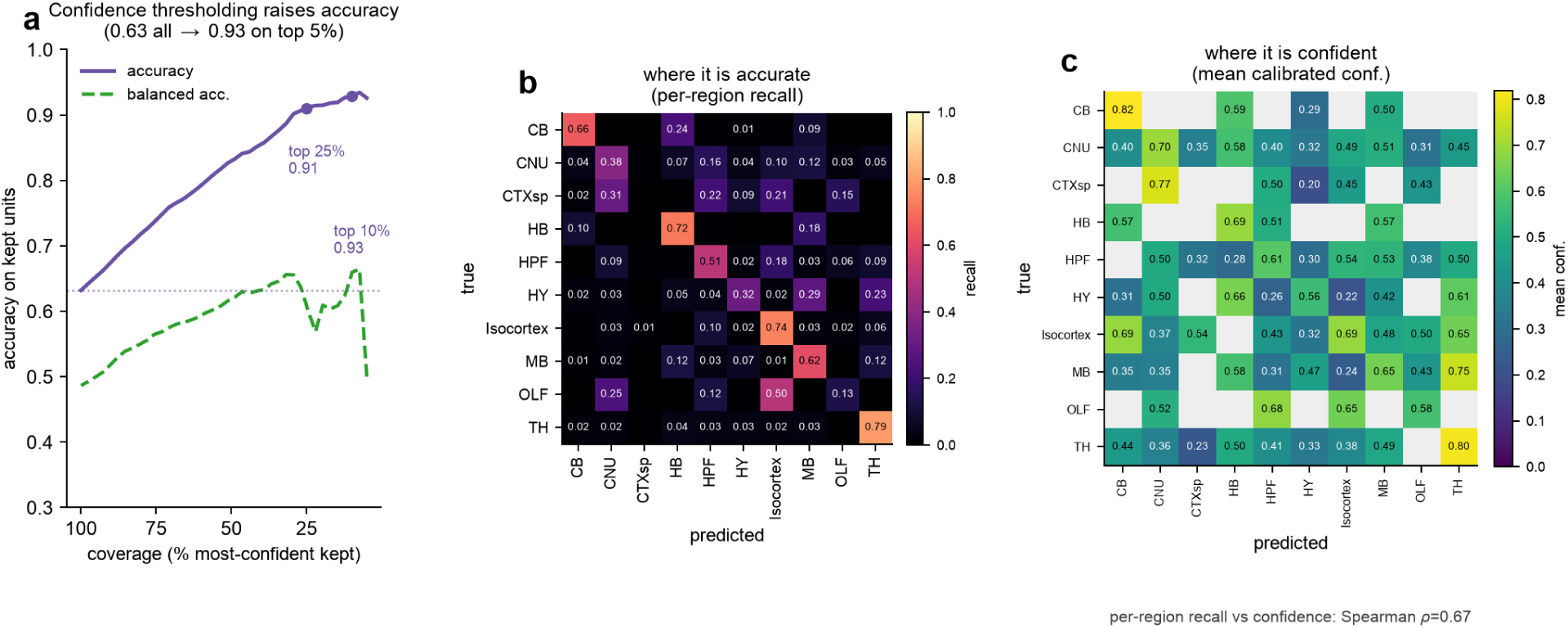
MYCEL’s calibrated confidence supports selective prediction on IBL Cosmos region decoding. **a** Selective prediction. Keeping only the most-confident predictions raises accuracy on the retained units from 0.63 over all units to 0.91 on the top 25%, plateauing near 0.93 on the top 10% and 5% by confidence (purple; dotted line, all-unit accuracy). Balanced accuracy (green dashed) climbs more gently, from 0.49 over all units to a peak near 0.66, because calibrated confidence correlates with how many units a region supplies, so high-confidence subsets are dominated by the well-sampled regions rather than balanced across them. **b**, Per-region recall (row-normalized confusion), showing where the model is accurate. **c**, Mean calibrated confidence in each (true, predicted) cell, showing where the model is confident (cells with *<* 3 predictions grayed). The two share the Cosmos-region order, and confidence tracks accuracy region by region; bright, high-recall diagonal entries are the high-confidence ones and the off-diagonal misclassifications are low-confidence (per-region recall versus diagonal confidence, Spearman *ρ* = 0.67). All panels use the temperature-scaled confidence, on the animal-disjoint test split.

**Supplementary Figure 17:**
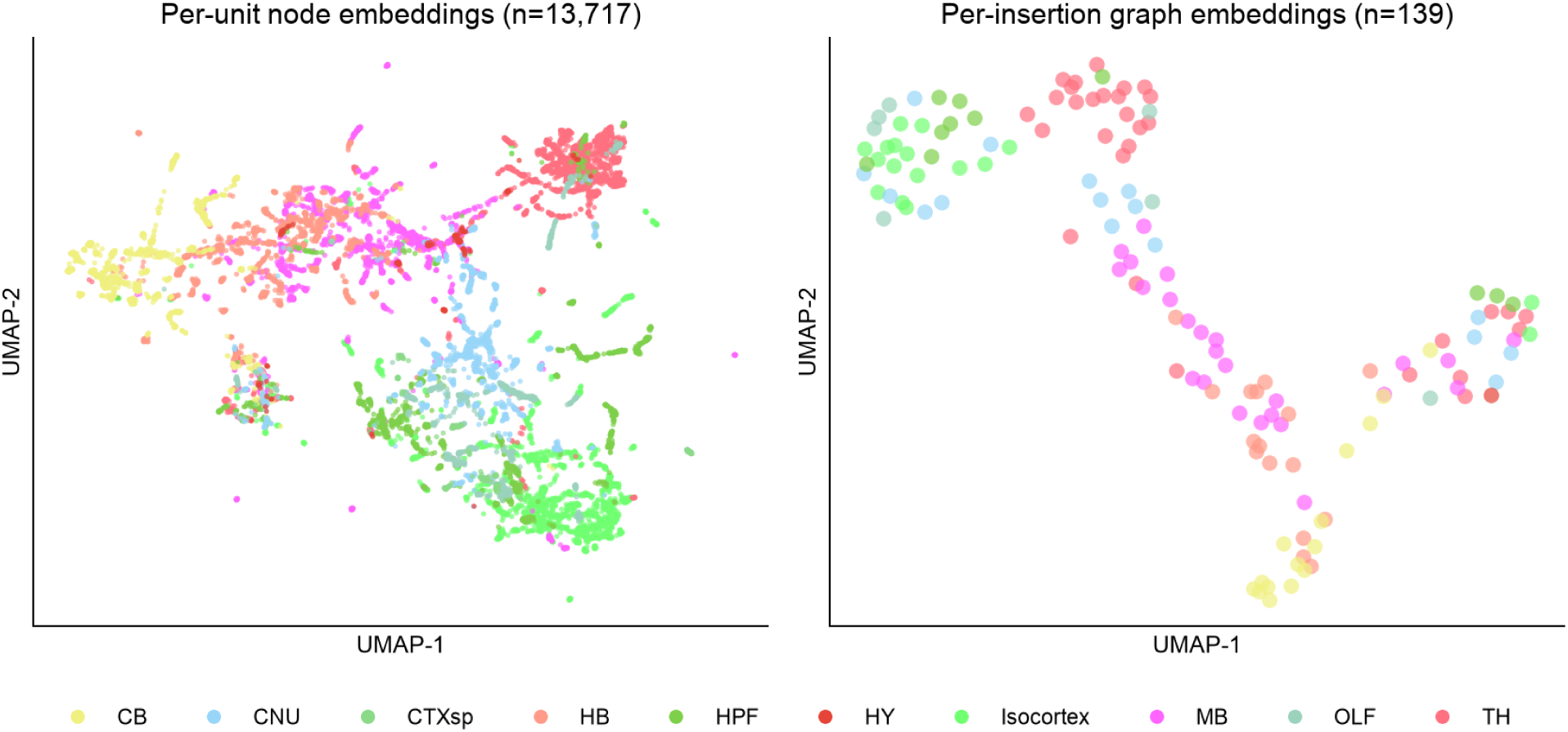
The region MYCEL’s learned embeddings organize by Cosmos region. UMAP of MYCEL’s learned embeddings on the held-out IBL good-units fold (fold 0), colored by the 10-region Cosmos taxonomy. **Left**, per-unit node embeddings after message passing (*n* = 13,717 held-out units); **right**, per-insertion graph-pooled embeddings (*n* = 139 insertions). Thalamus (TH), cerebellum (CB), isocortex and cerebral nuclei (CNU) occupy distinct territory, while the over-represented and adjacent regions (hippocampal formation HPF, olfactory areas OLF and midbrain MB) overlap (compare Supplementary Fig. 8). Cosmos colors follow the Allen ontology.

**Supplementary Figure 18:**
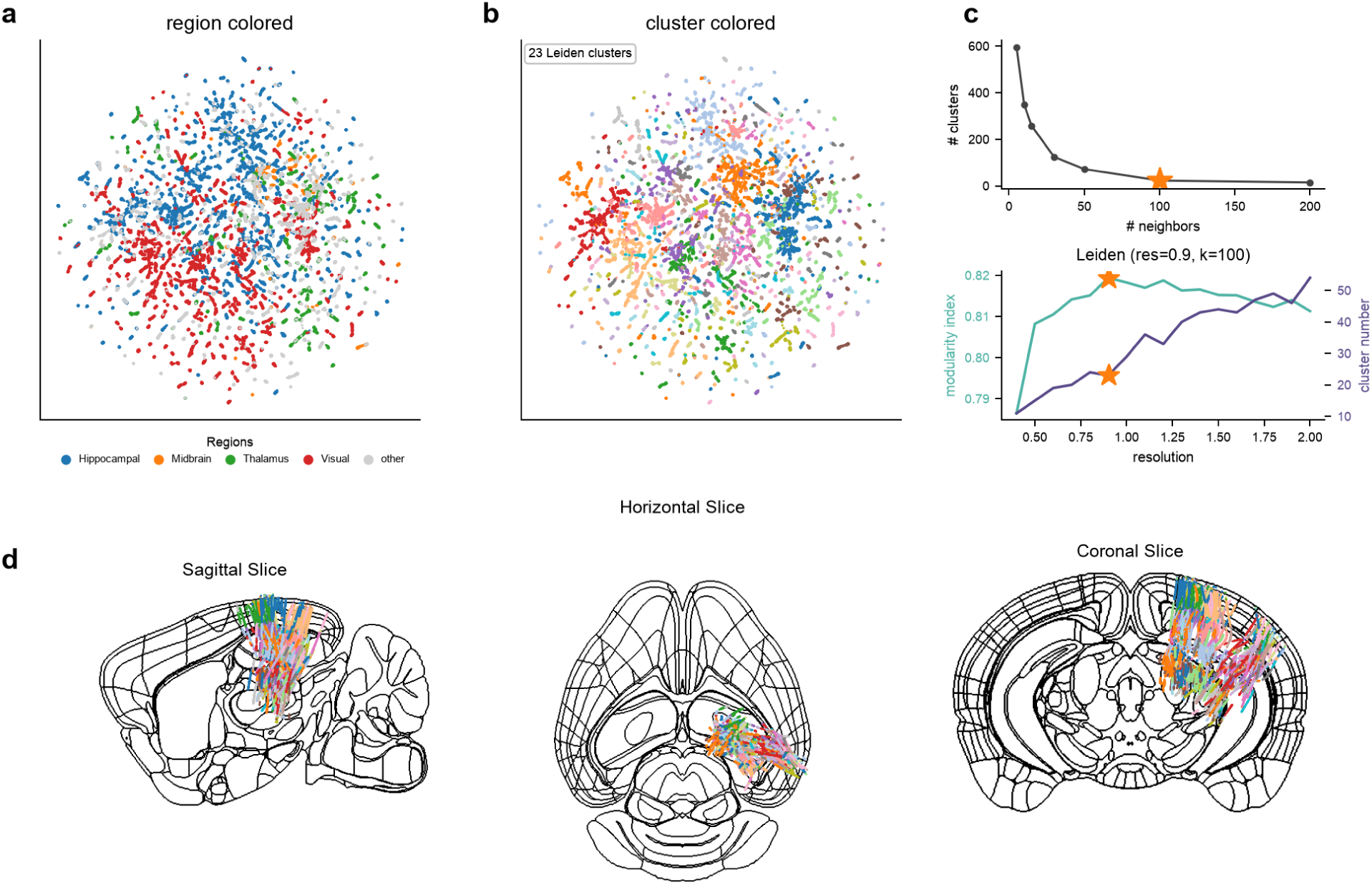
Unsupervised structure of the Allen region MYCEL embedding. As Fig. 3 but for the CCF-registered Allen Visual Coding corpus (*n* = 61,770 units). **a**, UMAP colored by verified Allen super-region (hippocampal, midbrain, thalamus, visual; gray, other structures); **b**, the same geometry colored by unsupervised Leiden community (23 communities); **c**, operating-point selection (neighbors = 100, resolution = 0.9 at maximum modularity); **d**, communities placed at their Allen CCF coordinates on the three boundary slices. Because the Allen units come from the area-subset probe set, they occupy a localized anatomical volume rather than the whole brain.

**Supplementary Figure 19:**
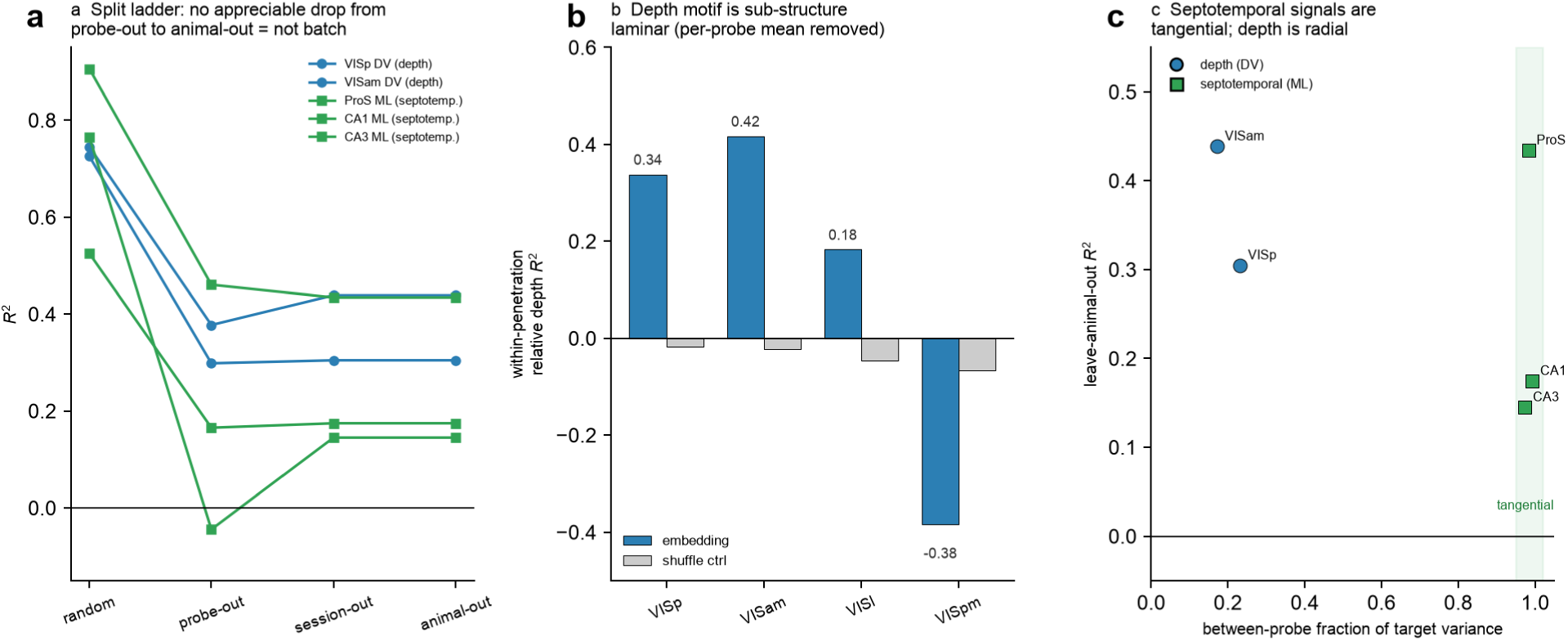
Controls for the two position motifs, on verified anatomy. The decode of Fig. 4 is stress-tested using each unit’s real probe, session and animal identity. **a**, Split series. For each motif signal, out-of-fold *R*^2^ under random-fold (leaky), leave-probe-out, leave-session-out and leave-animal-out. No signal falls appreciably from leave-probe-out to leave-animal-out (the drop from the leaky random fold is the removal of within-probe leakage), so the surviving value is localization, not session or animal batch. **b**, within-penetration relative depth (dorso-ventral position with each probe’s mean removed, leave-probe-out) versus a shuffle-probe negative control. After cross-probe placement is removed, the cortical depth motif remains within-structure and laminar (VISp *R*^2^ = 0.34, VISam 0.42), with the control near zero. **c**, Geometry of the two motifs, showing leave-animal-out *R*^2^ against the between-probe fraction of each target’s variance. The septotemporal (ML) signals sit at 97–99% between-probe, i.e. they are tangential to the penetration with almost no within-shank spread, unlike cortical depth, which is radial and varies along the penetration.

**Supplementary Figure 20:**
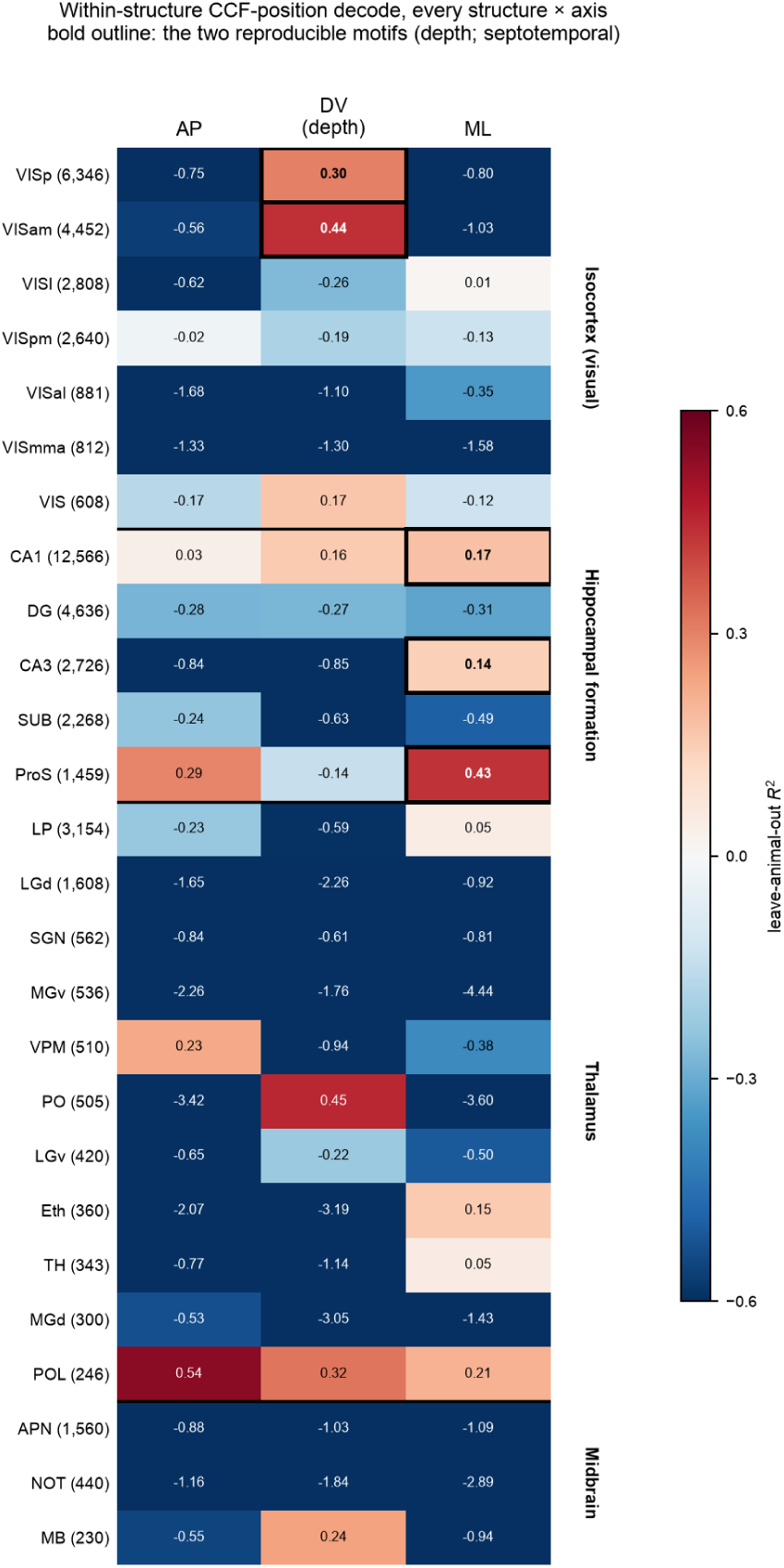
Within-structure CCF-position decode for every well-sampled structure. Leave-animal-out *R*^2^ for predicting each unit’s real Allen CCF coordinate (antero-posterior AP, dorso-ventral DV, medio-lateral ML) from its MYCEL node embedding, computed separately within each structure and grouped by super-region (rows). Verified anatomy (structure_acronym), real 3-axis CCF coordinates, ridge regression under a leave-animal-out split. Only structures with at least 200 units and 6 animals are shown, since leave-animal-out is unstable below this (unit count in parentheses). Color is *R*^2^ (diverging scale clipped at *±*0.6; the value is annotated in every cell). Bold outlines mark the two reproducible within-structure motifs quantified in Fig. 4 and Supplementary Fig. 19, cortical depth (VISp and VISam on DV) and hippocampal septotemporal position (ProS, CA1 and CA3 on ML). Isolated positive cells in small individual nuclei (for example PO or POL, with few hundred units and single-digit animal counts) are not reproducible across the split series and reflect the instability of leave-animal-out at low sample size, so they are not outlined.

**Supplementary Figure 21:**
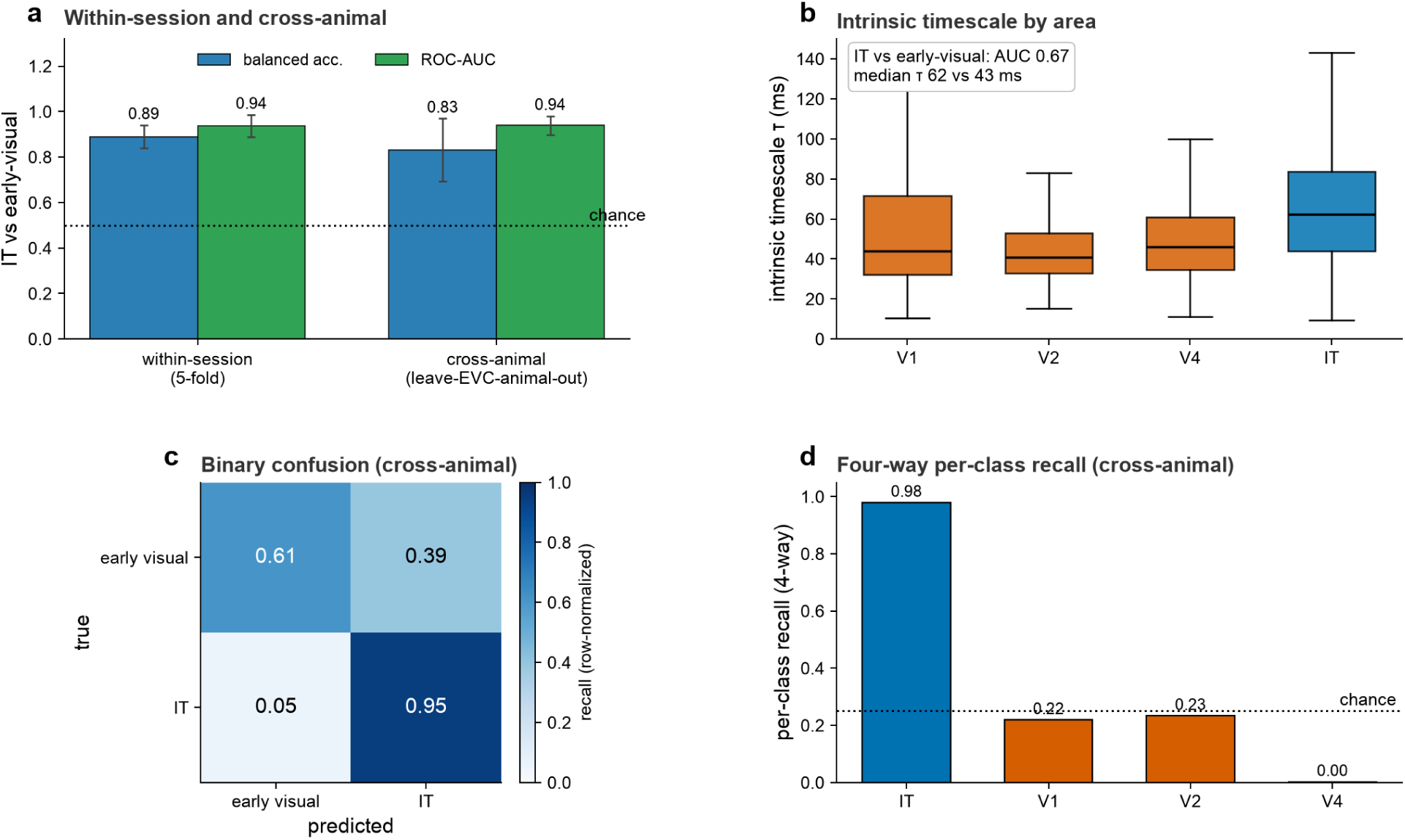
Macaque ventral stream cross-animal decode, intrinsic timescale and four-way partition. Two-class MYCEL (IT versus early/intermediate visual cortex, V1/V2/V4) on the Triple-N data, alongside Fig. 5a–d. **a**, Balanced accuracy and ROC-AUC within session (five folds) and across animals (leave-EVC-animal-out, two folds, read descriptively; Methods). **b**, Per-unit autocorrelogram intrinsic timescale *τ* by area (boxes, median and interquartile range (IQR); whiskers, 1.5*×*IQR; *n* = 14,600 fitted units); inset, IT versus EVC ROC-AUC and median *τ* (two-sided Mann–Whitney *P* = 1.6*×*10*^−^*^163^). **c**, Cross-animal confusion for the two-class decode (row-normalized recall). **d**, Per-class recall for the four-way V1/V2/V4/IT partition, across animals. In **a**, bars, mean; error bars, s.d. over folds. Dotted line, chance (0.5 two-class, 0.25 four-way).

**Supplementary Figure 22:**
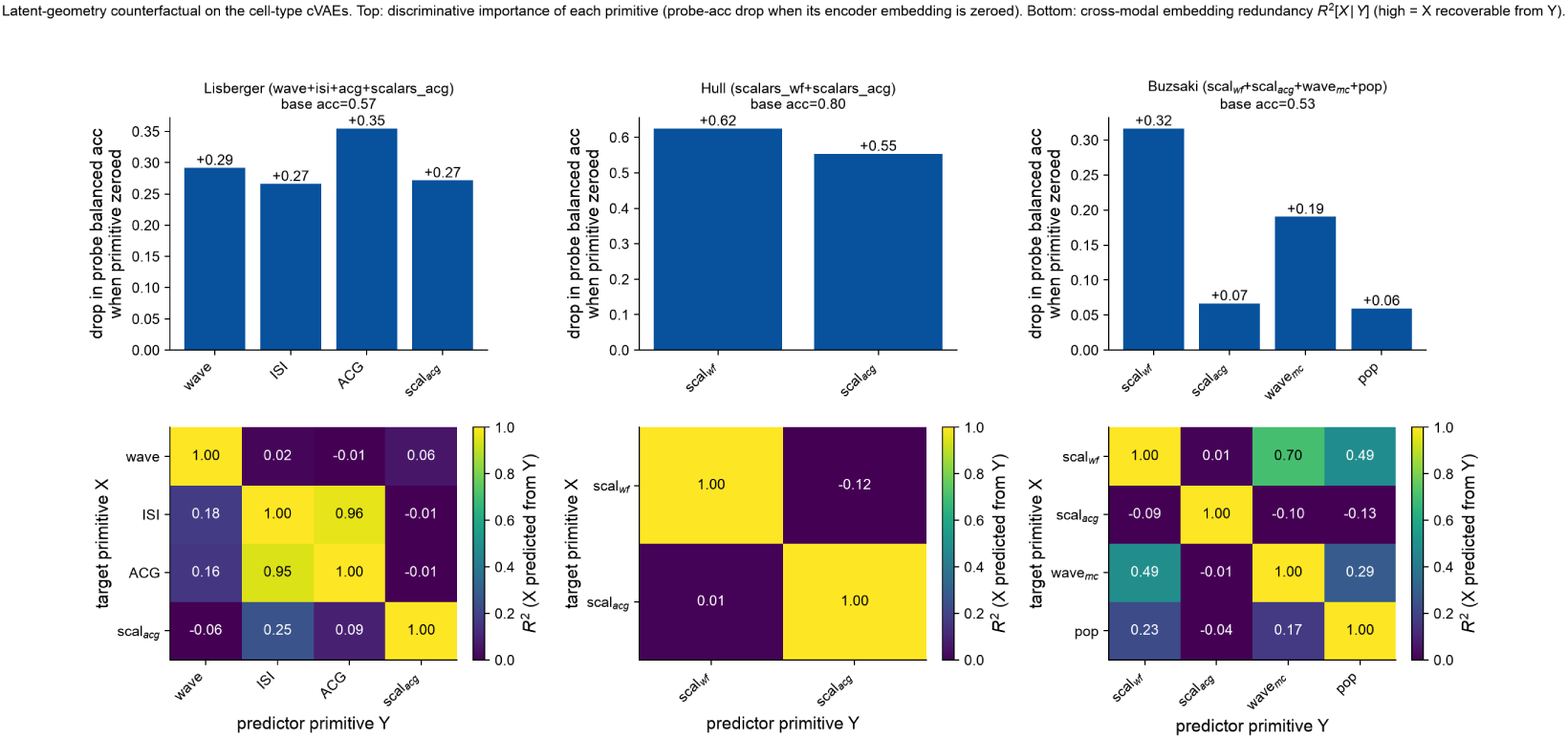
Latent-geometry counterfactual on the cell-type cVAEs. Hull and Lisberger use their search-selected cVAEs; Buzsáki uses a cVAE that includes the population primitive. Top, discriminative importance (drop in *k*-NN balanced accuracy when each primitive’s embedding is zeroed). Bottom, cross-modal embedding redundancy *R*^2^[*X | Y*]. Hull’s two scalar families are complementary and both critical; on Lisberger the waveform is unique and the added autocorrelogram-scalar block complementary; on Buzsáki the population primitive is the least important and only moderately unique.

**Supplementary Figure 23:**
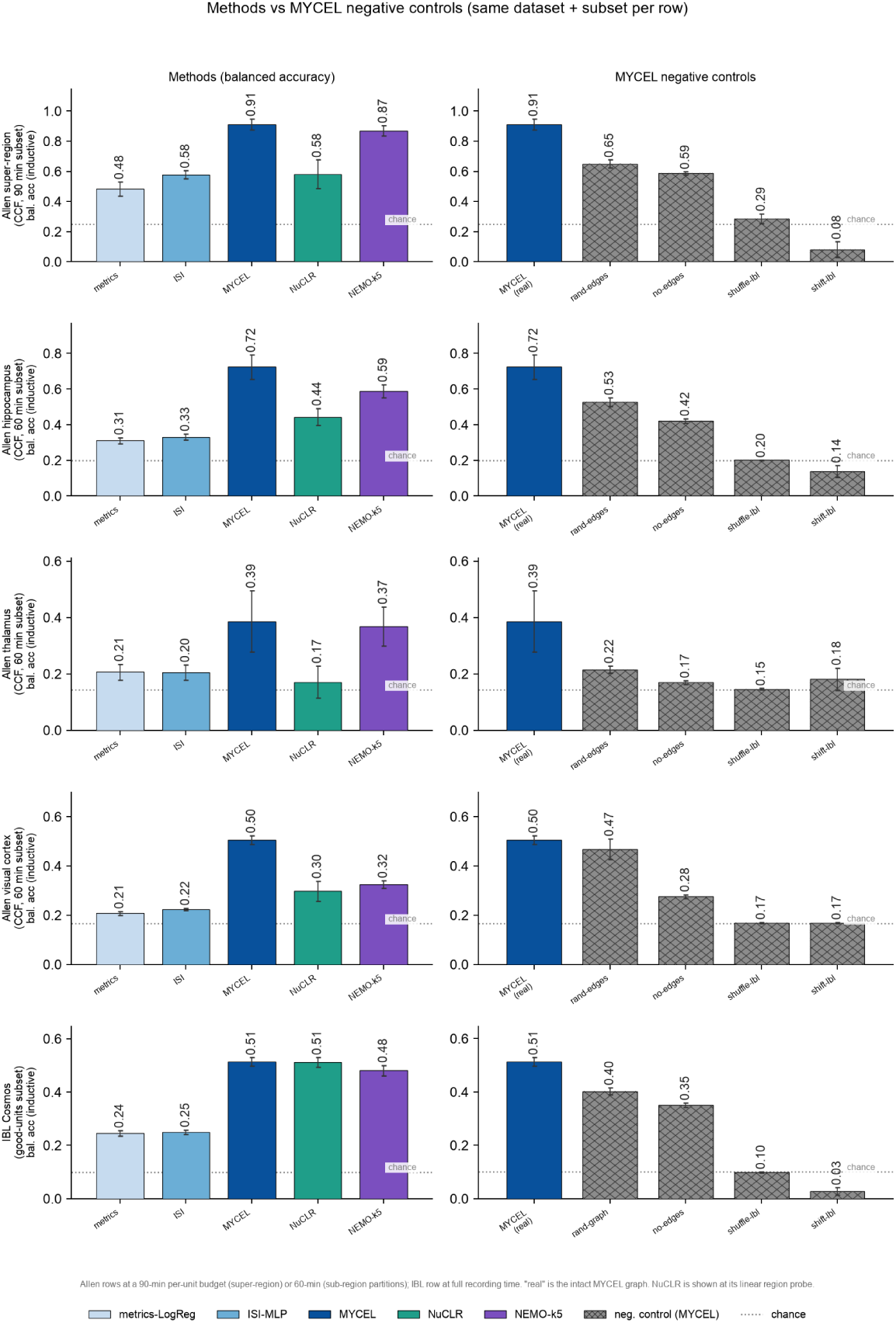
Method sweep and negative controls on every region task that carries both. Rows, the five region tasks for which both a method sweep and MYCEL negative controls were run, namely the Allen super-region (CCF-registered build, 90-min budget) and its hippocampal formation (5-way), thalamus (7-way) and visual areas (6-way) sub-region partitions (60-min budget), and the IBL Cosmos good-units task. **Left**, all methods on every row (single-unit metrics-LogReg and ISI-MLP; MYCEL; NuCLR; NEMO-k5). **Right**, MYCEL negative controls for the real graph, random edges, no edges, label-shuffle and label-shift. On the Allen super-region the real graph reaches *∼* 0.91, no-edges (node features only) still holds *∼* 0.59 and randomized edges *∼* 0.65, all far above the label-destruction floor (0.29 shuffle, 0.08 shift), so both the intrinsic node features and the graph topology carry regional signal that the other does not. MYCEL scores highest or ties on every task, ahead of both single-unit encoders (against its closest competitor among NuCLR and NEMO-k5, super-region 0.91 vs 0.87 at 90 min; hippocampal formation 0.72 vs 0.59 and visual areas 0.50 vs 0.32 at 60 min; the harder 7-way thalamus partition ties at *∼* 0.38 and IBL Cosmos ties NuCLR at *∼* 0.51), and clears its own controls wherever the task carries signal; on the thalamic partition the margin over the controls narrows. Bars, per-fold mean; whiskers, s.d.

**Supplementary Table 3:**
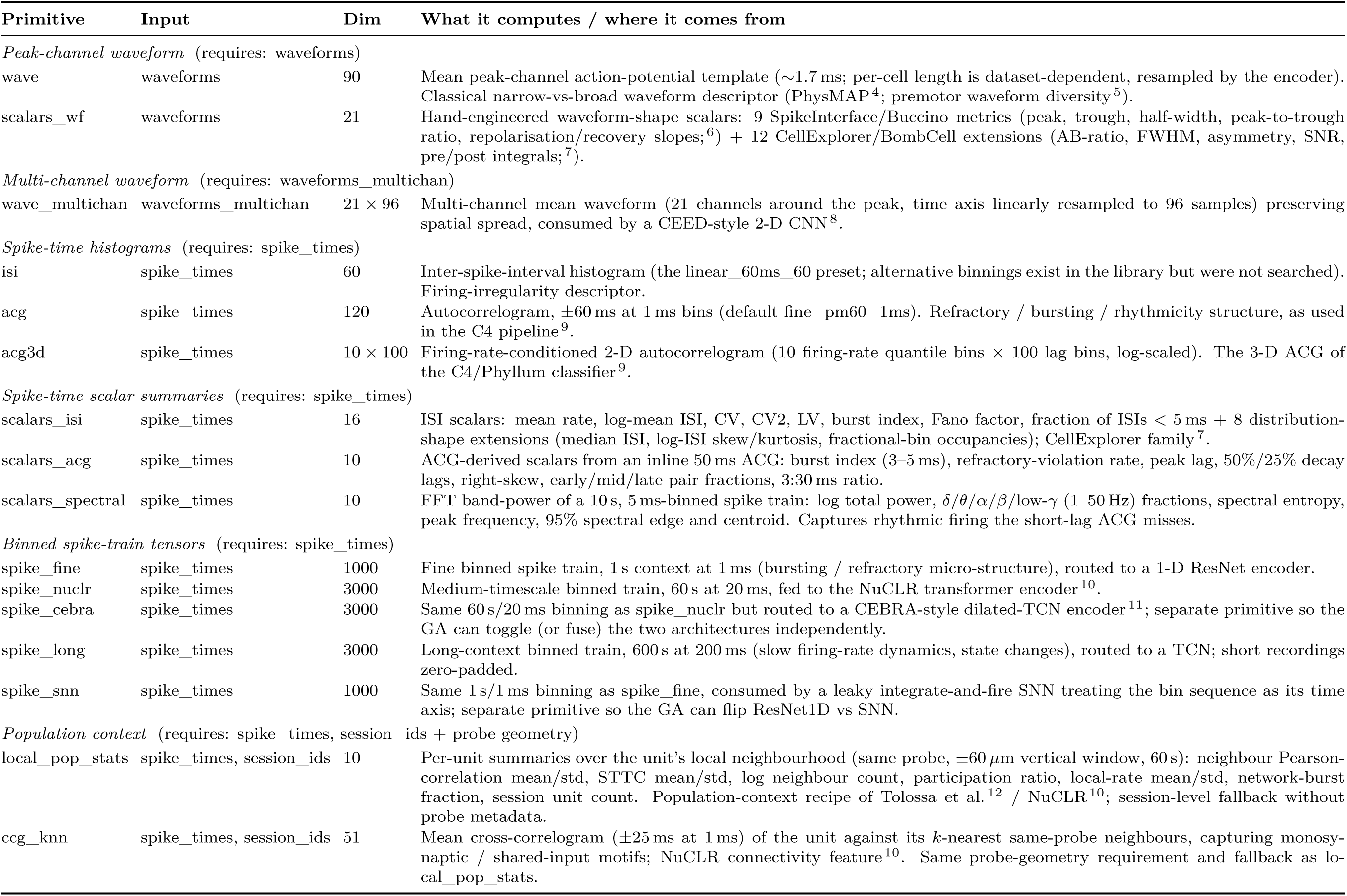
The 16 feature primitives evolved by the GA. Each primitive is one feature *modality* in the primitive library; it adapts a raw recording into a per-cell tensor consumed by a dedicated encoder. The GA evolves a binary on/off mask over these 16 primitives (plus the cVAE preset, latent width and *β*) with pymoo’s mixed-variable NSGA-II (nondominated sort then crowding distance), minimising four objectives: validation ELBO, *−*linear-probe balanced accuracy, number of active primitives, and log_10_ of an analytic complexity proxy (Methods). “Input” names the raw-recording field consumed; “Dim” is the per-cell output shape at the default binning preset. Rows are grouped by input/primitive type.

**Supplementary Table 4:** Resolved NSGA-II operator settings (pymoo v0.6.2 mixed-variable genetic algorithm; SBX, simulated binary crossover). Population 32, 16 generations, rank-and-crowding survival, random mating selection, duplicate elimination on.

| Setting (parameter type) | Crossover | Mutation | Notes |
| --- | --- | --- | --- |
| 16 primitive bits (binary) | uniform, $p_c=0.9$ | bit-flip, $p_m=0.0625$ | |
| $z_{\text{dim}}$ (integer, $[8, 48]$) | SBX, $\eta=15$ | polynomial, $\eta=20$ | rounding repair |
| $\beta$ (real, $[0.25, 4.0]$) | SBX, $\eta=15$ | polynomial, $\eta=20$ | |
| cVAE preset (categorical, 6) | uniform, $p_c=0.9$ | random choice | |

**Supplementary Table 5:** Class-weighted versus unweighted region decoding. (scalars_plus node features / depth edges; inverse-frequency weights capped 10*×*; inductive-animal, mean over 5 matched splits, chance 0.10). Weighting trades plain accuracy for balanced accuracy and rare-region recall (e.g. IBL HY recall 0.07 *→* 0.55) while leaving macro-F1 essentially unchanged: the per-region recall deficit is an imbalance effect, not a feature ceiling.

| Method / unit set | Acc. (unwt.) | bAcc. (unwt.) | Acc. (wt.) | bAcc. (wt.) |
| --- | --- | --- | --- | --- |
| MYCEL, IBL good | 0.67 | 0.51 | 0.61 | 0.53 |
| MYCEL, IBL all | 0.75 | 0.58 | 0.70 | 0.60 |

